# Mapping neuropeptide signaling in the human brain

**DOI:** 10.1101/2024.12.11.627947

**Authors:** Eric G. Ceballos, Asa Farahani, Zhen-Qi Liu, Filip Milisav, Justine Y. Hansen, Alain Dagher, Bratislav Misic

**Affiliations:** Montréal Neurological Institute, McGill University, Montréal, QC, Canada

## Abstract

Neuropeptides are functionally diverse signaling molecules in the brain, regulating a wide range of basal bodily and cognitive processes. Despite their importance, the distribution and function of neuropeptides in the human brain remains underexplored. Here we comprehensively map the organization of human whole-brain neuropeptide receptors across multiple levels of description; from molecular and cellular embedding to mesoscale connectivity and macroscale cognitive specialization. Using gene transcription as a proxy, we reconstruct a topographic cortical and subcortical atlas of neuropeptide receptors for 38 neuropeptide receptors, across 14 different neuropeptide families. We find that most neuropeptide receptors are highly expressed either in cortex or subcortex, delineating an anatomical cortical-subcortical gradient. Neuropeptides preferentially co-localize with metabotropic neurotransmitters, suggesting a system-wide correspondence between slow-acting molecular signaling mechanisms. Mapping neuropeptide receptors and their cognate ligands onto white-matter connectomes, we demonstrate that specific neuropeptides families shape electrophysiological and haemodynamic interregional connectivity. To investigate the behavioural consequences of distributed neuropeptide systems, we apply meta-analytic decoding to neuropeptide maps and show a gradient of functions, from sensory-cognitive to reward and bodily functions. Finally, evolutionary analysis indicates extended positive selection for neuropeptides in early mammals, suggesting that refinement of neuropeptides coincides with the emergence of neocortex and higher cognitive function. Collectively, these results show that the neuropeptide receptors are highly organized across the human brain and closely intertwined with multiple features of brain structure and function.

## INTRODUCTION

The brain is organized across multiple spatial scales, from microscale molecular gradients to mesoscale connections and interactions between neuronal populations to macroscale morphological patterns. While these organizational features can be measured and mapped using different techniques, how they link together remains an important question. A dominant paradigm has been to study interactions among neural elements. For instance, multiple studies report evidence that local microarchitectural features shape global connectomic wiring [12], such as gene transcription [53, 133, 139], cell morphology[62, 148], laminar differentiation [17, 18, 63, 173, 181] and intracortical myelin [13, 17, 53, 77, 133, 148]. Of particular interest are signaling molecules that support communication across multiple scales, such as neurotransmitters and their receptors [52, 67, 151].

Neuropeptides are a fundamental molecular signaling system both in the nervous system and the body. Like metabotropic neurotransmitters, neuropeptides support inter-cellular communication via G protein-coupled receptors. Compared to neurotransmitters, neuropeptides are larger amino acid chains that are released from presynaptic axonal terminals into extra-cellular space, thereby acting over a wide range of neighbouring post-synaptic boutons. Neuropeptides are thought to be synthesized and secreted in a small set of structures, such as the hypothalamus, although there is increasing evidence that both neuropeptides and their receptors are considerably more widespread throughout the brain and body [134, 192]. Importantly, neuropeptides support a rich repertoire of functions, including sleep (e.g., hypocretin/orexin)[92, 125], pain (e.g., nociceptin/orphanin) [57, 166], feeding (e.g., leptin) [10, 121], reward (e.g., opioids) [31, 57] and social cognition (e.g., oxytocin) [9, 93, 129, 146]. As a result, neuropeptides mediate how the brain perceives and manipulates both internal bodily and external environments. Despite their importance, how neuropeptide signaling maps onto the organization of the brain remains poorly understood.

Here we comprehensively map neuropeptide systems in the human brain. Using gene transcription as a proxy, we chart the spatial distribution of 38 neuropeptide receptors across 14 different neuropeptide families in both cortex and subcortex. We first study how neuropeptide systems co-localize with other canonical structural and functional features of the brain. We then annotate neural wiring patterns with neuropeptide distributions to understand how neuropeptide signaling contributes to connectome organization. Finally, we trace the phylogenetic emergence of neuropeptide systems in the brain and map these systems onto patterns of regional functional specialization. Altogether, we demonstrate that the spatial patterning of neuropeptide receptors is highly organized across the brain and has multiple functional consequences.

## RESULTS

A whole-brain atlas of neuropeptide receptors was assembled for 38 neuropeptide receptors across 14 different neuropeptide families (Fig. 1). Topographic receptor maps were reconstructed using the intensity of the corresponding microarray gene transcript from the Allen Human Brain Atlas (AHBA) as a proxy [71]. Receptors were mapped to the Schaefer 400 cortical atlas [147], the Melbourne Subcortex Atlas S4 [167], and the hypothalamic delineation from the CIT168 atlas [118], yielding a whole-brain atlas with 455 regions. Briefly, gene transcripts were filtered based on multiple quality control criteria, including mean intensity, correlation with RNA-seq and differential stability (Table 1; see *Methods*). To ensure specificity of findings, we compare results with sets of null genes, pseudorandomly selected such that they have comparable magnitude distribution and spatial autocorrelation with the transcript markers for each receptor (see *Methods*). To validate the spatial distribution of receptor gene expression in AHBA, we systematically compare it with an independent RNA-seq atlas (Human Protein Atlas; Fig. S1) [155, 170].

**TABLE 1.** Neuropeptide receptor overview. Neuropeptide receptor distributions were estimated using gene expression from the Allen Human Brain Atlas [70, 71]. Gene transcripts were filtered based on multiple quality control criteria, including (1) mean intensity > 0.2, (2) correlation with RNA-seq > 0.2, and (3) differential stability > 0.1. For reference, mean expression within cortex, hypothalamus and subcortex are shown.

| Gene | Full name | Family | Quality control |  |  | Mean gene expression |  |  |
| --- | --- | --- | --- | --- | --- | --- | --- | --- |
|  |  |  | Mean intensity | Corr. RNA-seq | Diff. stability | Cortex | Hypo-thalamus | Subcortex |
| ADCYAP1R1 | ADCYAP receptor type I | Glucagon/secretin | 0.97 | 0.45 | 0.35 | 0.35 | 0.81 | 0.64 |
| ADIPOR2 | Adiponectin receptor 2 | Adipose | 0.95 | 0.54 | 0.14 | 0.52 | 0.42 | 0.60 |
| APLNR | Apelin receptor | Unclassified | 0.75 | 0.62 | 0.30 | 0.49 | 0.75 | 0.56 |
| BDKRB2 | Bradykinin receptor B2 | Kinin/tensin | 0.39 | 0.66 | 0.40 | 0.45 | 0.11 | 0.32 |
| CALCRL | Calcitonin receptor like receptor | Calcitonin | 0.79 | 0.39 | 0.18 | 0.47 | 0.46 | 0.44 |
| CCKBR | Cholecystokinin B receptor | CCK/gastrin | 0.61 | 0.66 | 0.54 | 0.87 | 0.16 | 0.28 |
| EDNRA | Endothelin receptor type A | Endothelin | 0.42 | 0.60 | 0.32 | 0.53 | 0.32 | 0.41 |
| EDNRB | Endothelin receptor type B | Endothelin | 0.96 | 0.78 | 0.54 | 0.40 | 0.86 | 0.81 |
| GALR1 | Galanin receptor 1 | Galanin | 0.37 | 0.48 | 0.38 | 0.53 | 0.68 | 0.37 |
| GHR | Growth hormone receptor | Unclassified | 0.63 | 0.70 | 0.44 | 0.42 | 0.69 | 0.54 |
| GIPR | Gastric inhibitory polypeptide receptor | Glucagon/secretin | 0.97 | 0.20 | 0.15 | 0.39 | 0.51 | 0.56 |
| GLP2R | Glucagon like peptide 2 receptor | Glucagon/secretin | 0.31 | 0.73 | 0.59 | 0.67 | 0.28 | 0.26 |
| GRPR | Gastrin releasing peptide receptor | Bombesin-like | 0.28 | 0.25 | 0.27 | 0.54 | 0.45 | 0.21 |
| HCRTR1 | Hypocretin receptor 1 | Unclassified | 0.38 | 0.58 | 0.18 | 0.48 | 0.65 | 0.50 |
| LEPR | Leptin receptor | Adipose | 0.76 | 0.20 | 0.26 | 0.37 | 0.72 | 0.64 |
| MCHR1 | Melanin concentrating hormone receptor 1 | Unclassified | 0.34 | 0.58 | 0.35 | 0.60 | 0.19 | 0.17 |
| MCHR2 | Melanin concentrating hormone receptor 2 | Unclassified | 0.45 | 0.75 | 0.42 | 0.60 | 0.09 | 0.10 |
| NPFFR2 | Neuropeptide FF receptor 2 | F-/Y-amide | 0.51 | 0.42 | 0.36 | 0.49 | 0.49 | 0.18 |
| NPR2 | Natriuretic peptide receptor 2 | Natriuretic factor | 0.99 | 0.23 | 0.19 | 0.45 | 0.70 | 0.51 |
| NPR3 | Natriuretic peptide receptor 3 | Natriuretic factor | 0.25 | 0.42 | 0.34 | 0.50 | 0.53 | 0.31 |
| NPY1R | Neuropeptide Y receptor Y1 | F-/Y-amide | 0.69 | 0.80 | 0.66 | 0.60 | 0.55 | 0.45 |
| NPY2R | Neuropeptide Y receptor Y2 | F-/Y-amide | 0.29 | 0.60 | 0.16 | 0.47 | 0.42 | 0.21 |
| NPY5R | Neuropeptide Y receptor Y5 | F-/Y-amide | 0.53 | 0.45 | 0.41 | 0.56 | 0.54 | 0.60 |
| NTSR1 | Neurotensin receptor 1 | Kinin/tensin | 0.22 | 0.81 | 0.67 | 0.46 | 0.48 | 0.31 |
| NTSR2 | Neurotensin receptor 2 | Kinin/tensin | 0.98 | 0.83 | 0.44 | 0.44 | 0.89 | 0.73 |
| OPRK1 | Opioid receptor kappa 1 | Opioid | 0.43 | 0.71 | 0.54 | 0.48 | 0.53 | 0.59 |
| OPRL1 | Opioid related nociceptin receptor 1 | Opioid | 0.90 | 0.65 | 0.15 | 0.65 | 0.57 | 0.44 |
| OPRM1 | Opioid receptor mu 1 | Opioid | 0.60 | 0.55 | 0.62 | 0.38 | 0.66 | 0.58 |
| OXTR | Oxytocin receptor | Vasopressin/oxytocin | 0.68 | 0.61 | 0.44 | 0.38 | 0.93 | 0.66 |
| RAMP1 | Receptor activity modifying protein 1 | Calcitonin | 1.00 | 0.50 | 0.21 | 0.46 | 0.70 | 0.72 |
| RAMP3 | Receptor activity modifying protein 3 | Calcitonin | 0.72 | 0.68 | 0.51 | 0.50 | 0.56 | 0.36 |
| RXFP1 | Relaxin/insulin like family peptide receptor 1 | Insulin | 0.52 | 0.67 | 0.51 | 0.74 | 0.12 | 0.17 |
| SORT1 | Sortilin 1 | Kinin/tensin | 1.00 | 0.46 | 0.34 | 0.55 | 0.16 | 0.43 |
| SSTR1 | Somatostatin receptor 1 | Somatostatin | 0.50 | 0.86 | 0.75 | 0.59 | 0.66 | 0.28 |
| SSTR2 | Somatostatin receptor 2 | Somatostatin | 0.98 | 0.80 | 0.49 | 0.66 | 0.31 | 0.35 |
| TACR2 | Tachykinin receptor 2 | Kinin/tensin | 0.71 | 0.29 | 0.18 | 0.49 | 0.39 | 0.37 |
| VIPR1 | Vasoactive intestinal peptide receptor 1 | Glucagon/secretin | 0.40 | 0.47 | 0.35 | 0.82 | 0.15 | 0.32 |
| VIPR2 | Vasoactive intestinal peptide receptor 2 | Glucagon/secretin | 0.51 | 0.85 | 0.58 | 0.67 | 0.29 | 0.27 |

**Figure 1.**
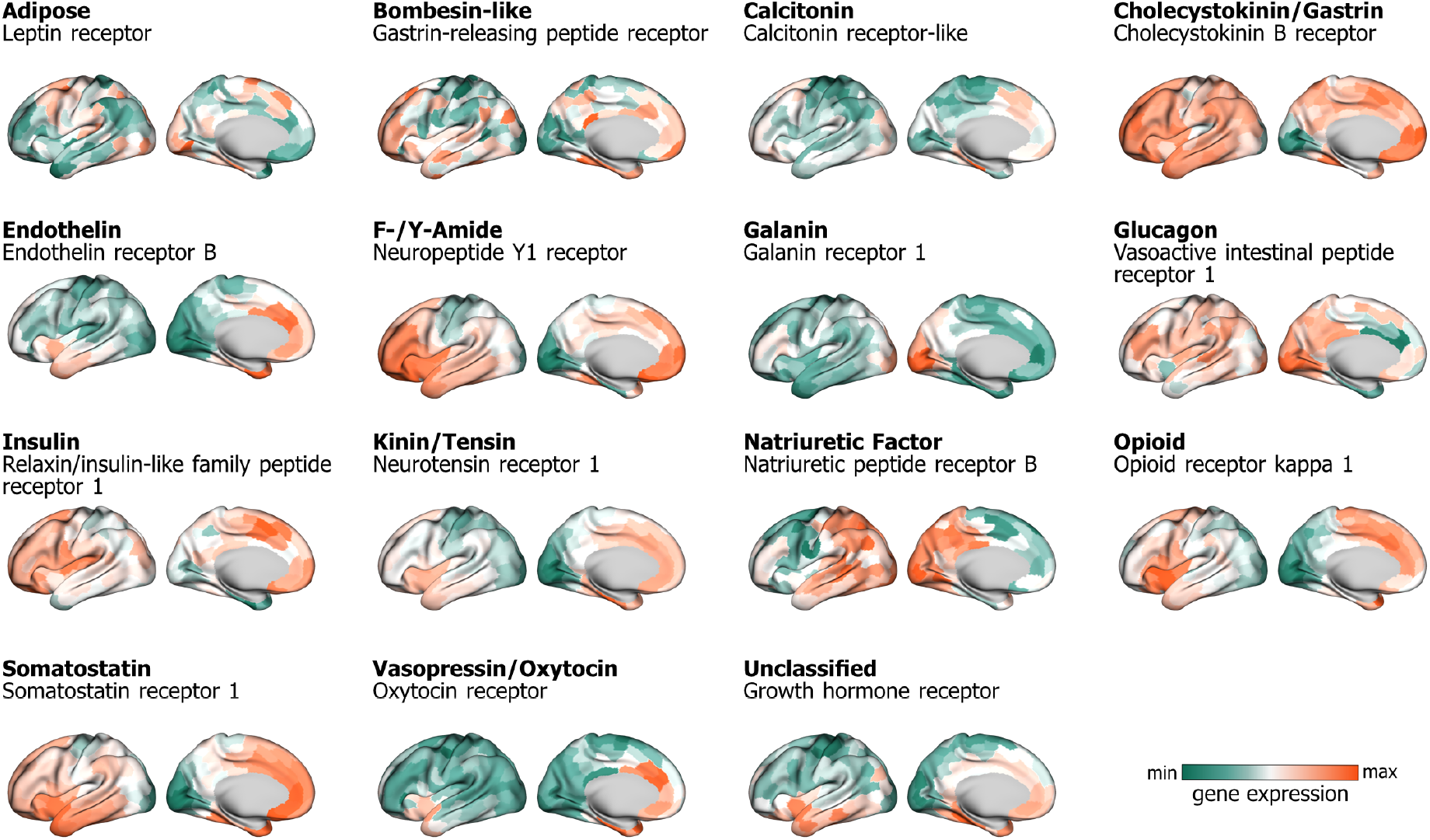
Neuropeptide families. We map 38 neuropeptide receptors based on their gene expression available in the Allen Human Brain Atlas [71]. Out of 38 neuropeptide receptors, we identify 14 families of neuropeptides from an established taxonomy of peptides [21]. For each neuropeptide family (bold font), a single exemplar receptor is shown (regular font). Maps are min-max scaled. For simplicity, only cortical topographies are shown, but note that all analyses reported were carried out for both cortex and subcortex. Technical details about gene expression quality control metrics and mean expression in specific structures are shown in Table 1.

### Mapping neuropeptide signaling in the human brain

Fig. 1 shows the cortical spatial distribution for one exemplar receptor in each neuropeptide family. The spatial distributions display considerable heterogeneity and concordance with prior literature. For example, oxytocin receptors are highly expressed in limbic cortex, consistent with their role in social cognition [9, 93, 129]. Importantly, there is substantial diversity of receptor expression, motivating a further exploration of how neuropeptide receptors are embedded within structural and functional organization of the brain.

To more comprehensively describe the anatomical distribution of neuropeptide receptors, we estimate the mean expression of receptor genes in cortex (stratified by intrinsic networks), hypothalamus and subcortex (Fig. 2a). The receptors are organized in a cortical-subcortical axis, with some receptors preferentially expressed in cortex, e.g., VIP receptors, and others expressed in hypothalamus and other subcortical nuclei. Fig. 2b summarizes this gradient of expression, showing the median expression of each neuropeptide receptor within cortex, hypothalamus and subcortex. For example, we observe high expression of oxytocin receptors in amygdala, opioid receptors in the nucleus accumbens, and the leptin receptor in hypothalamus and hippocampus. Interestingly, there are numerous instances of neuropeptide receptors that are primarily associated with basal bodily functions but that are nevertheless highly expressed in the brain, and in cortex in particular (e.g. cholecystokinin B receptor). Much like vasoactive intestinal peptide (VIP) receptors, which are now known to be expressed in inhibitory cortical interneurons [123, 124, 142], these examples contribute to a growing realization that, despite their names and initially-ascribed functions, neuropeptide receptors can be expressed in the brain and may potentially have additional functions in neural communication [134]. Note that several commonly studied neuropeptides did not pass quality control criteria and were therefore not considered, such as corticotropin releasing hormone (CRH) and ghrelin (see Table S1).

**Figure 2.**
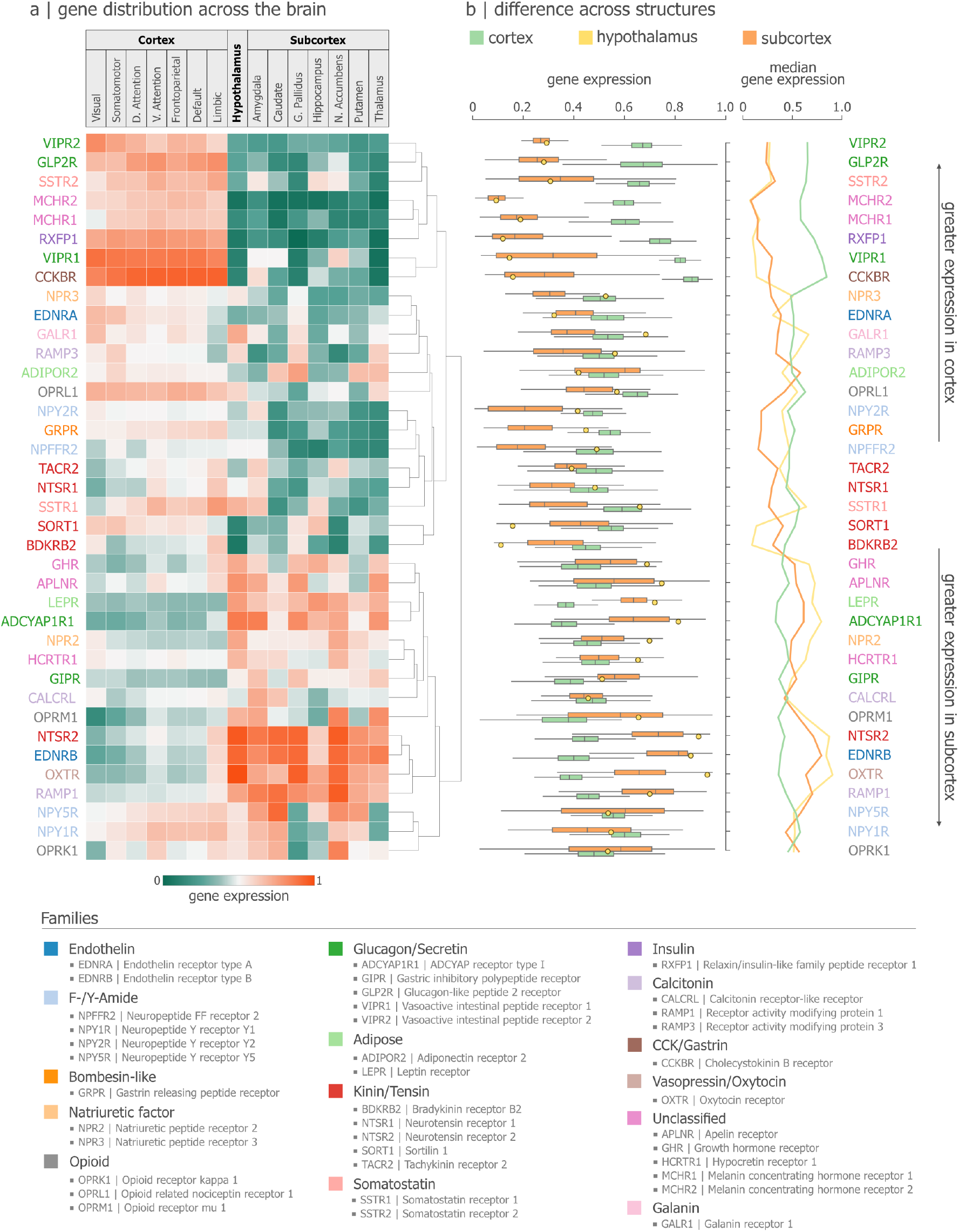
Neuropeptide distribution across the human brain. Neuropeptide receptor topographies stratified according to anatomical and functional landmarks. (a) Mean expression of individual neuropeptide receptors in cortex, hypothalamus and subcortex. Cortical receptor expression is stratified according to the canonical intrinsic functional networks (Visual, Somatomotor, Dorsal Attention, Ventral Attention, Frontoparietal, Default Mode, and Limbic) [147]. Subcortical receptor expression is stratified according to a functional atlas (Amygdala, Caudate Nucleus, Globus Pallidus, Hippocampus, Nucleus Accumbens, Putamen, and Thalamus) [167]. Hypothalamic expression was localized using the CIT168 atlas [118]. (b) Boxplots of neuropeptide receptor expression within cortex, subcortex and hypothalamus, with medians plotted on the right side. Receptors are coloured according to family membership, and vertically sorted using hierarchical clustering.

### Co-localization with neurotransmitter receptors

Just like neurotransmitters, neuropeptides are chemical messengers that work by binding to G protein-coupled (metabotropic) receptors, modulating the excitability of proximal cells. The primary difference is that neurotransmitter release is more directed towards the postsynaptic terminal, whereas neuropeptides are secreted more tangentially and diffusely, also binding to receptors located on neighbouring cells (Fig. 3a). Neurotransmitters and neuropeptides therefore occupy the same local environment, naturally raising the question of how these two systems converge spatially.

**Figure 3.**
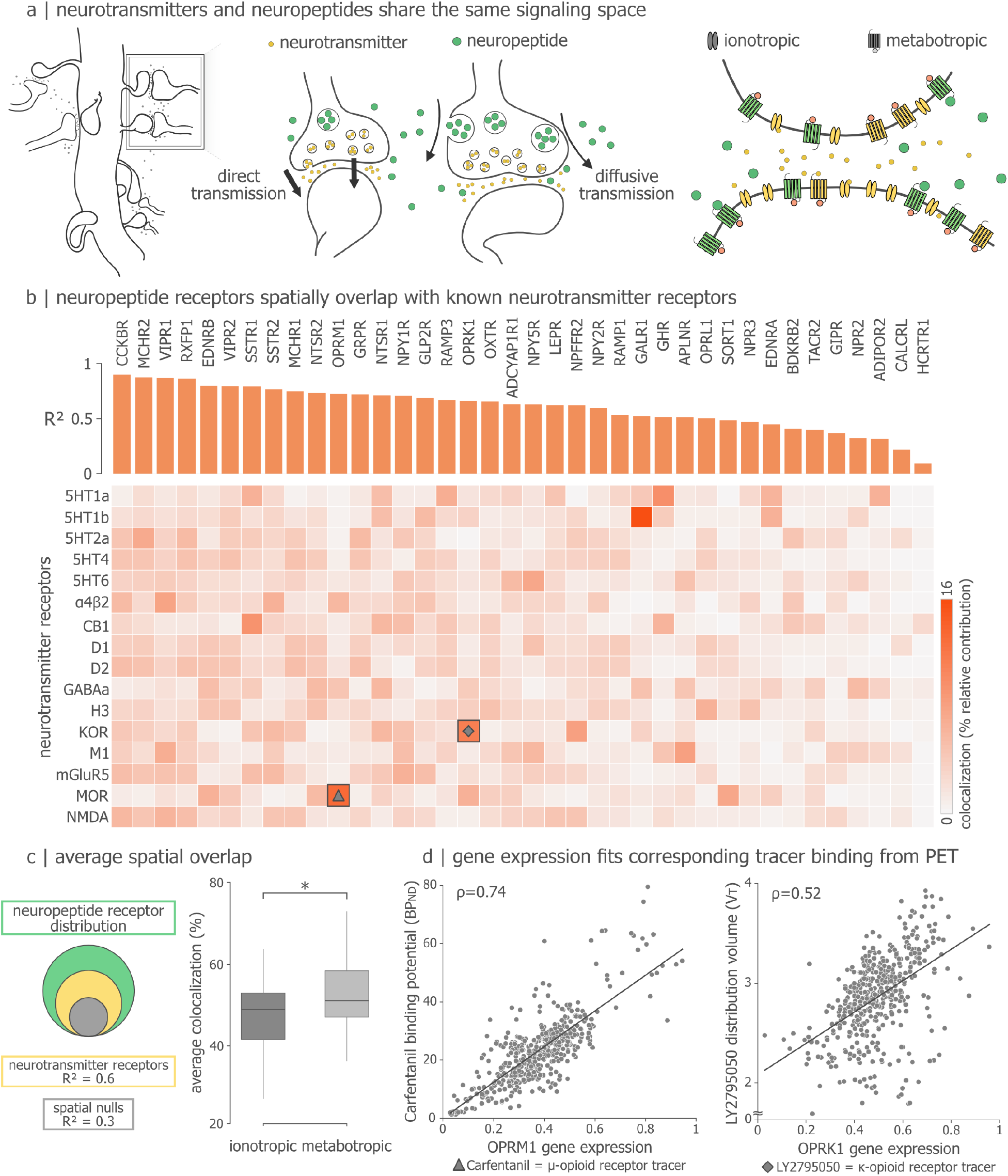
Co-localization with neurotransmitter receptors. (a) Schematic showing different types of synaptic chemical signaling, including direct ionotropic and metabotropic neurotransmission and diffusive neuropeptide signaling. (b) Neuropeptide receptor distributions are correlated with 16 neurotransmitter receptor densities estimated using PET imaging [67]. Vertical bars indicate the adjusted *R*^2^ when predicting each neuropeptide receptor map using a linear combination of neurotransmitter receptor maps. Heatmap colour intensity corresponds to the relative contribution of each neurotransmitter receptor map to the spatial distribution of a neuropeptide receptor [6]. (c) Mean co-localization (*R*^2^) between neuropeptide and neurotransmitter receptors. (d) Scatterplots show the spatial correspondence between gene expression-estimated and PET imaging-estimated receptor densities of the *µ*- and *κ*-opioid receptors. *µ*- and *κ*-opioid receptors were imaged using [11C]Carfentanil and [11C]LY2795050 radiotracers, respectively.

Here we compare the spatial patterns of neuropeptide receptors with spatial patterns of 16 neurotransmitter receptors, derived from a whole-brain positron emission tomography (PET) atlas [67]. Notably, the atlas contains both metabotropic and fast-acting ionotropic receptors (*α*4*β*2, M1, NMDA, GABAa). Fig. 3b shows how well the spatial distribution of each neuropeptide receptor can be predicted as a linear combination of neurotransmitter receptor densities. Neuropeptide receptors are ordered according to their adjusted *R*^2^ (orange bars), and the heatmap shows the percent contribution (i.e. colocalization) of each neurotransmitter receptor [6] (see *Methods*). On average, neurotransmitter receptor distributions share 60% spatial variance with neuropeptide receptors (Fig. 3c). When stratified into ionotropic and metabotropic receptors, we find that metabotropic receptors are significantly more co-localized with neuropeptide receptors (*t*(74) = 2.71, *p* = 0.007; Fig.3c), suggesting that neuropeptide receptors are prevalent in the synaptic environment where metabotropic neurotransmitter receptors reside.

The analysis above also offers an opportunity to conduct a validation analysis. Namely, the PET atlas contains two tracers ([11C] LY2795050 and [11C] Carfentanil) that measure the receptor protein density of two opioid receptors that are part of our gene library: *κ*-opioid receptor (KOR) and *µ*-opioid receptor (MOR). To assess whether neuropeptide gene expression is an acceptable proxy for the corresponding protein, we focus on those two entries in the Fig. 3a heatmap, indicated by diamond and triangle symbols. Fig. 3d shows that the spatial correlation between the genes (OPRK1 and OPRM1) and PET-estimated protein densities is positive and statistically significant in both cases (KOR: *ρ* = 0.52, *p*_SMASH_ = 0.0015; MOR: *ρ* = 0.74, *p*_SMASH_ = 0.0006), offering reassurance that neuropeptide distributions can be inferred from gene expression (see *Discussion*).

### Neuropeptide signaling in whole-brain connectivity

So far, we focused on where neuropeptide receptors are situated within the anatomical organization of the brain. Now, we consider how neuropeptide signaling supports inter-regional communication. We first ask where the signaling neuropeptide molecules (as opposed to receptors) are expressed, such as neuropeptide Y, leptin, calcitonin and galanin. Stratifying the expression of 41 different neuropeptide precursor genes according to their anatomical location, we find the greatest expression in hypothalamus (*t* = 2.04, *p* = 0.02) and cortex (*t* = 1.86, *p* = 0.03; Fig. 4a) compared to subcortex, consistent with the notion that the hypothalamus is the nexus for neuropeptide signaling [74, 101, 138, 183]. Pursuing the idea that neuropeptide signaling is aligned with hypothalamic function, we correlate the spatial distribution of neuropeptide receptors across cortical and subcortical brain regions with the magnitude of resting state fMRI functional connectivity to hypothalamus (Fig. 4b). Contrasting this relationship to random genes with matched spatial autocorrelation and value distribution, we find that a subset of peptide receptors are aligned to the hypothalamic connectivity pattern beyond the null distribution. These receptor families – tensin, endothelin, opioid, Y-amide, somatostatin, galanin – are putatively part of a larger signaling infrastructure built around hypothalamic function.

**Figure 4.**
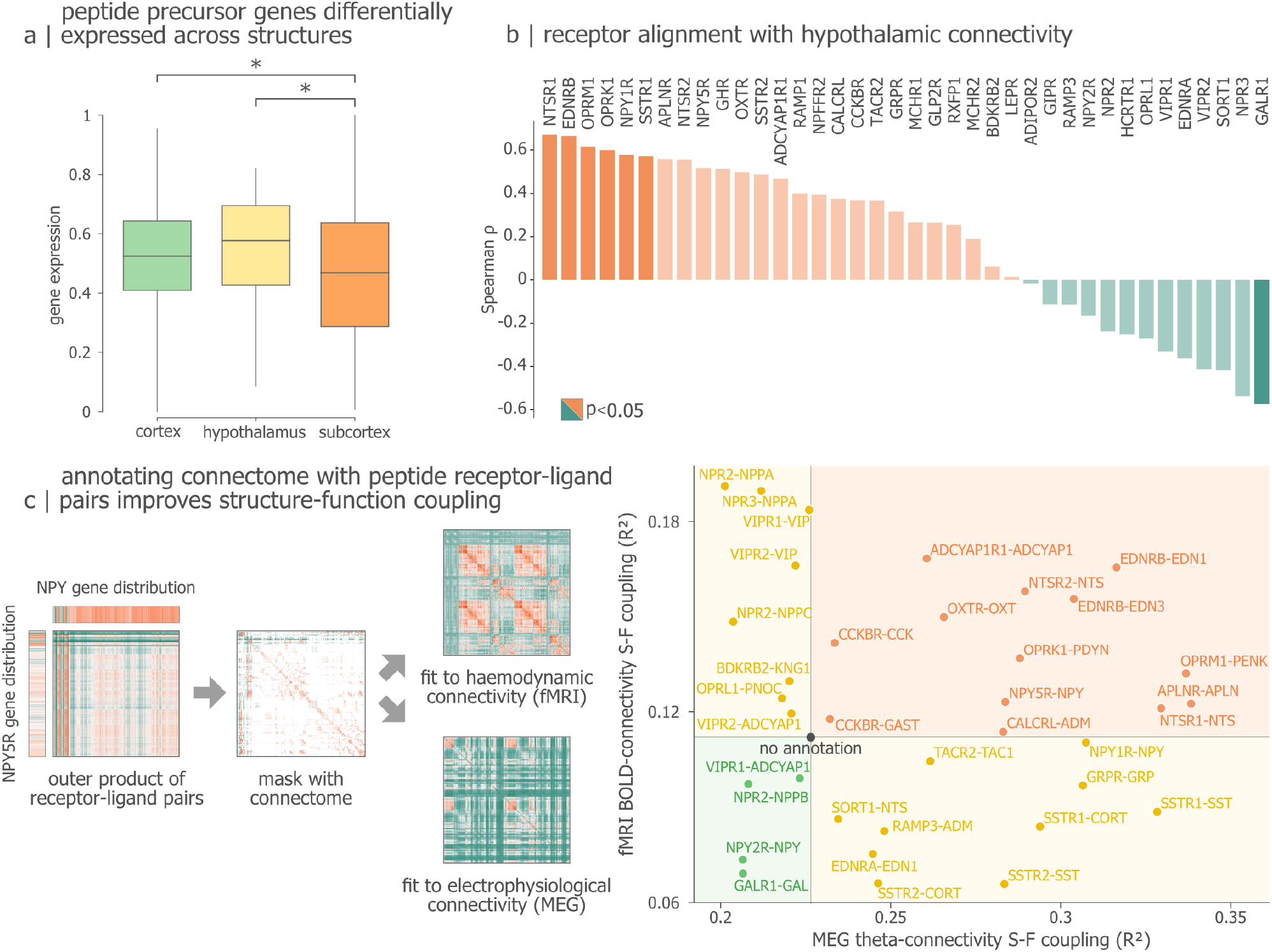
Peptide signaling boosts structure-function coupling. (a) Mean expression of neuropeptide ligand precursor genes in cortex, hypothalamus and subcortex. (b) Whole-brain maps of hypothalamic seed connectivity were estimated using resting state fMRI functional connectivity [27, 154]. Bars show the spatial correlations between hypothalamic connectivity maps and neuropeptide receptor maps. Significance of correlations was assessed using (1) gene nulls with matching spatial autocorrelation and value distribution, and (2) spatial autocorrelation-preserving nulls [24]. (c) Neuropeptide signaling networks were estimated by multiplying receptor-ligand densities between regions and subsequently masking with a template structural connectome derived using diffusion-weighted MRI. Functional connectivity estimated using resting state fMRI (haemodynamic connectivity) as well as source-resolved resting state magnetoencephalography (MEG; electrophysiological connectivity). Models of structure-function coupling were then fitted using structural connectivity only (“no annotation”), or using structural connectivity plus neuropeptide annotations. The magnitude of structure-function coupling is shown separately for each neuropeptide receptor-ligand pair, and for each functional connectivity mode. The figure shows MEG connectivity in the theta band (5-7 Hz). Results for other canonical electrophysiological frequency bands, including *δ* (2-4 Hz), *α* (8-14 Hz), *β* (15-29 Hz), low *γ* (30-59 Hz) and high *γ* (60-90 Hz) are shown in Fig. S6.

Extending the scope of inquiry beyond the hypothalamus, we finally ask to what extent does neuropeptide signaling contribute to global communication in the brain? To address this question, we construct a series of simple models in which we predict fMRI and MEG functional connectivity from dMRI structural connectivity enriched with neuropeptide annotations. Briefly, we construct a receptor-ligand matrix for each peptide by multiplying the magnitude of regional receptor expression with the expression of the corresponding ligand [70, 134, 157]. This matrix is then masked by a white matter connectome reconstructed using dMRI tractography such that it retains only edges that correspond to anatomical projections (Fig. 4c). The resulting neuropeptide signaling matrix is then compared with multiple functional connectivity modes estimated using resting-state fMRI-BOLD (haemodynamic connectivity) and resting-state MEG (*δ*-, *θ*-, *α*-, *β*-, and *γ*-electrophysiological connectivity) [67, 94, 149, 177, 187]. This comparison results in a measure of how structure-function coupling benefits from incorporating the location of neuropeptide receptors and their cognate ligands into structural connectome weights.

Fig. 4c shows the performance of annotated connectomes at fitting functional connectivity. To assess the increase in structure-function coupling brought by annotating the connectome with receptor-ligand pairs, we benchmark our results to a baseline result where no annotation is used, i.e., communication dynamics are modelled from the binary connectome template. We observe that specific pairings excel more in matching one mode of functional connectivity above the other. Neuropeptides from the natriuretic family (NPR2-NPPA, NPR3-NPPA & NPR2-NPPC) bias structure-function coupling towards haemodynamic connectivity. Natriuretic peptides are known to modulate blood pressure [109], offering an explanation for the positive steer towards functional connectivity that originates from neurovascular coupling. Receptor-ligand pairs from the opioid (OPRM1-PENK & OPRK1-PDYN), endothelin (EDNRB-EDN1 & EDNRB-EDN2) and tensin (NTSR1-NTS) families bias structure-function coupling positively towards both BOLD- and theta-connectivity. This is in line with previous reports of such peptides modulating signaling [2, 26, 28, 32, 58, 136] and neurovascular [61, 110, 188] activity. Moreover, pairings in the somatostatin (SSTR1-SST & SSTR1-CORT) and Y-amide (NPY1R-NPY) families show below baseline correspondence with BOLD-connectivity, but excel at aligning structural communication to theta-oscillatory activity. Again, our results fall in line with previous studies reporting the involvement of such peptides in theta-modulations [76, 168]. More importantly, the difference in performance between modes of connectivity demonstrates that neuropeptide signaling coincides with different communication pathways. This is underscored with the performance of galanin (GALR1-GAL), which does not enrich structural communication towards BOLD- or theta-connectivity, but is informative of low and high gamma-connectivity (see Fig. S6 for an extensive benchmark of neuropeptide enrichment in other bands such as *δ*-, *θ*-, *α*-, *β*-, and *γ*-bands). It is noteworthy that many of the highest-performing pairs in this analysis were also highlighted as participating in hypothalamic connectivity (Fig. 4a), suggesting that neuropeptide signaling originating from the hypothalamus shapes multiple modes of global communication in the brain.

### Neuropeptides delineate cognitive domains

In the previous subsections we explored principles that govern the regional distributions of neuropeptide receptors and the coordination of inter-regional neuropeptide signaling. We next investigate how neuropeptides shape cognitive specialization. To wit, initial discoveries about neuropeptide functions were based on behavioural changes in response to pharmacological manipulation, such as sleep and hypocretin [92], or weight-regulation and leptin [191]. Here we sought to map regional neuropeptide receptor distributions to patterns of regional cognitive specialization using the Neurosynth meta-analytic engine [186]. Briefly, Neurosynth is a meta-analytic tool that identifies regional activations associated with specific cognitive functions and psychological processes by aggregating data from thousands of fMRI studies. We cross-reference Neurosynth terms with terms from the Cognitive Atlas to derive 125 unique meta-analytic maps [126]. The resulting maps represent the likelihood that specific brain regions are associated with a given task, e.g., “reading” or “eating”, thereby serving as a reverse-inference engine from terms to neural activity.

To derive a data-driven mapping between receptors and terms, we use an unsupervised multivariate pattern learning technique, partial least squares correlation (PLSC) [102]. Briefly, PLSC seeks to identify weighted linear combinations of receptors and terms that optimally covary with each other across the brain [66]. We identify a single dominant mapping between receptors and terms (81.17% covariance accounted for, mean out-of-sample correlation = 0.36; Fig. 5a), and test it against (1) 10 000 parametric spatial autocorrelation preserving null sets (*p* = 0.0001) [23, 100], and (2) 10 000 sets of random genes with matching spatial autocorrelation and value distribution (see *Methods*; *p* = 0.0001). This major common axis displays a division between cortex and subcortex as evidenced by the disjoint distributions of scores. This latent variable is reminiscent of the cortex-subcortex gradient of neuropeptide receptors observed earlier (Fig. 2), revealing the cognitive architecture that mirrors this chemoarchitectural gradient.

**Figure 5.**
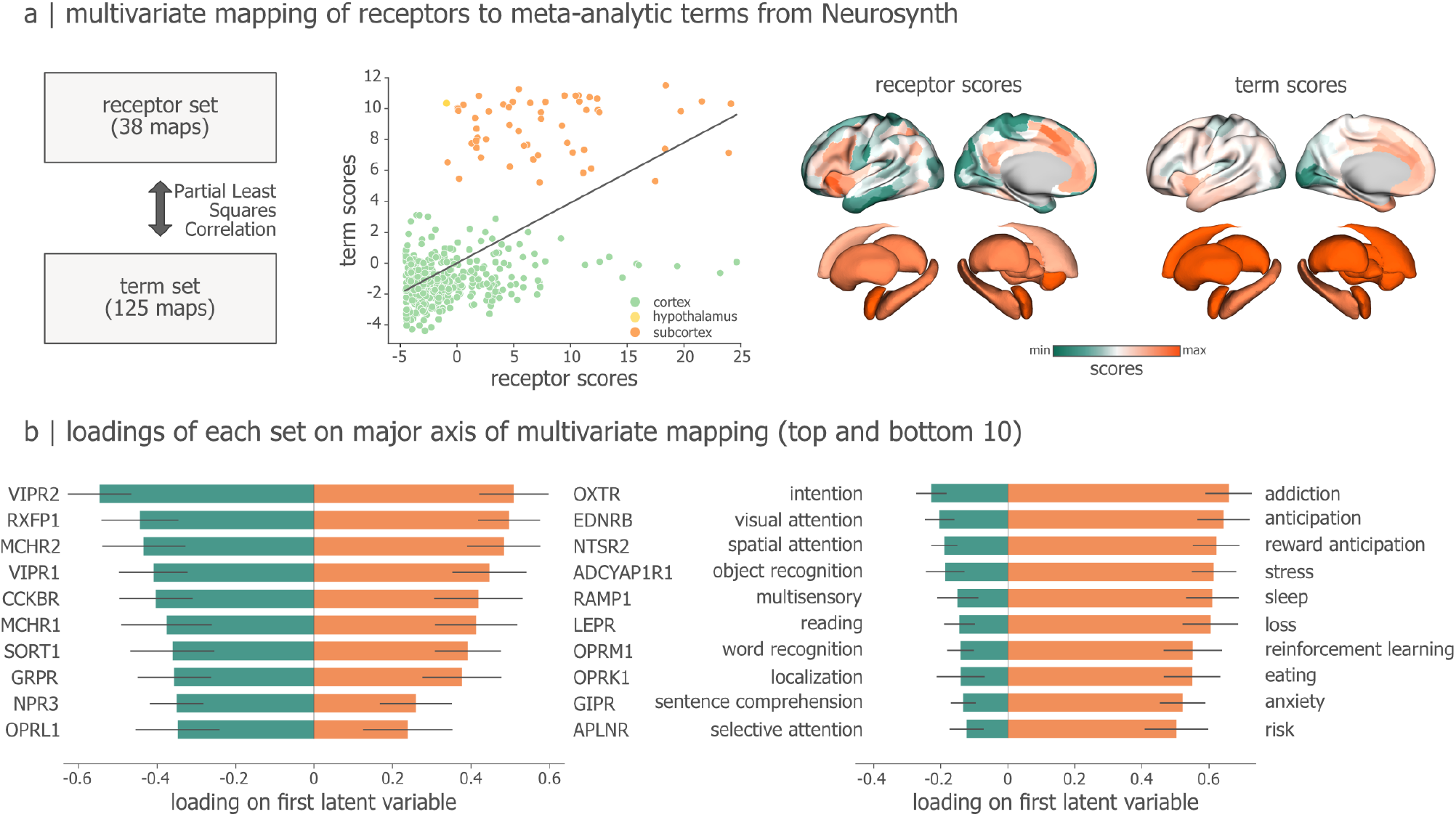
Neuropeptide receptor alignment with cognitive domains. (a) Partial least squares correlation was used to map the 38 neuropeptide receptor maps to a set of 125 cognitive terms from the Neurosynth meta-analytic atlas [186]. The analysis identified one significant latent variable. Term and receptor scores are shown as both a scatter plot and surface renderings. (b) Receptor and term loadings for the first latent variable. Error bars represent bootstrap-estimated 95% confidence intervals.

To understand the specific receptor-term mappings that constitute this gradient, we inspect the receptor and term loadings, i.e., the degree with which receptors and terms are correlated with the latent variable (Fig. 5b). On the negative side, we observe that cortically expressed neuropeptide receptors like VIPR1/2, MCHR1/2, RXFP1, CCKBR (negative bars on the left) covary with sensory processing terms such as “visual/spatial attention”, “object recognition”, “reading”, or “localization”. On the positive side, we observe that more subcortically expressed receptors (OXTR, EDNRB, NTSR2, ADCYAP1R1, RAMP1, LEPR, OPRM1) covary with terms such as “addiction”, “anticipation”, “risk”. These receptors and terms are prominent in the reward and reinforcement learning literature [29, 57, 84, 90, 128, 182], which involves the mesolimbic (predominantly subcortical) reward circuit [29, 112, 143, 184]. Additionally, we find positive loadings for terms related to emotional regulation such as “stress”, “loss”, or “anxiety”, as well as basal bodily functions such as “eating” or “sleep”. Collectively, these results show that neuropeptides are highly anatomically organized and support a spectrum of basal bodily to higher cognitive functions. This differentiated function of neuropeptides may potentially reflect homeostatic and allostatic demands rooted in environmental demands and evolutionary adaptation, a question that we pursue in the next subsection.

### Evolution of molecular signaling systems

To investigate the evolutionary roots of neuropeptide signaling, we conclude by conducting an exploratory evolutionary analysis focusing on a set of twelve species preceding the Homo sapiens species. These species were selected to represent key points along the vertebrate lineage [129, 146], which are: Petromyzon marinus (sea lamprey), Carcharodon carcharias (great white shark), Danio rerio (zebrafish), Xenopus tropicalis (western clawed frog), Gallus gallus (chicken), Ornithorhynchus anatinus (platypus), Sarcophilus harrisii (Tasmanian devil), Dasypus novemcinctus (nine-banded armadillo), Bos taurus (cattle), Mus musculus (house mouse), Macaca mulatta (rhesus macaque), and Pan troglodytes (chimpanzee) (Fig. 6a). The species relatedness among vertebrates is reflected in the gradually increasing amino acid similarity of neuropeptide receptors (Fig. 6b). However, while molecular similarity identifies shared ancestry [119], it does not identify the specific points in evolution at which adaptive changes occurred [85].

**Figure 6.**
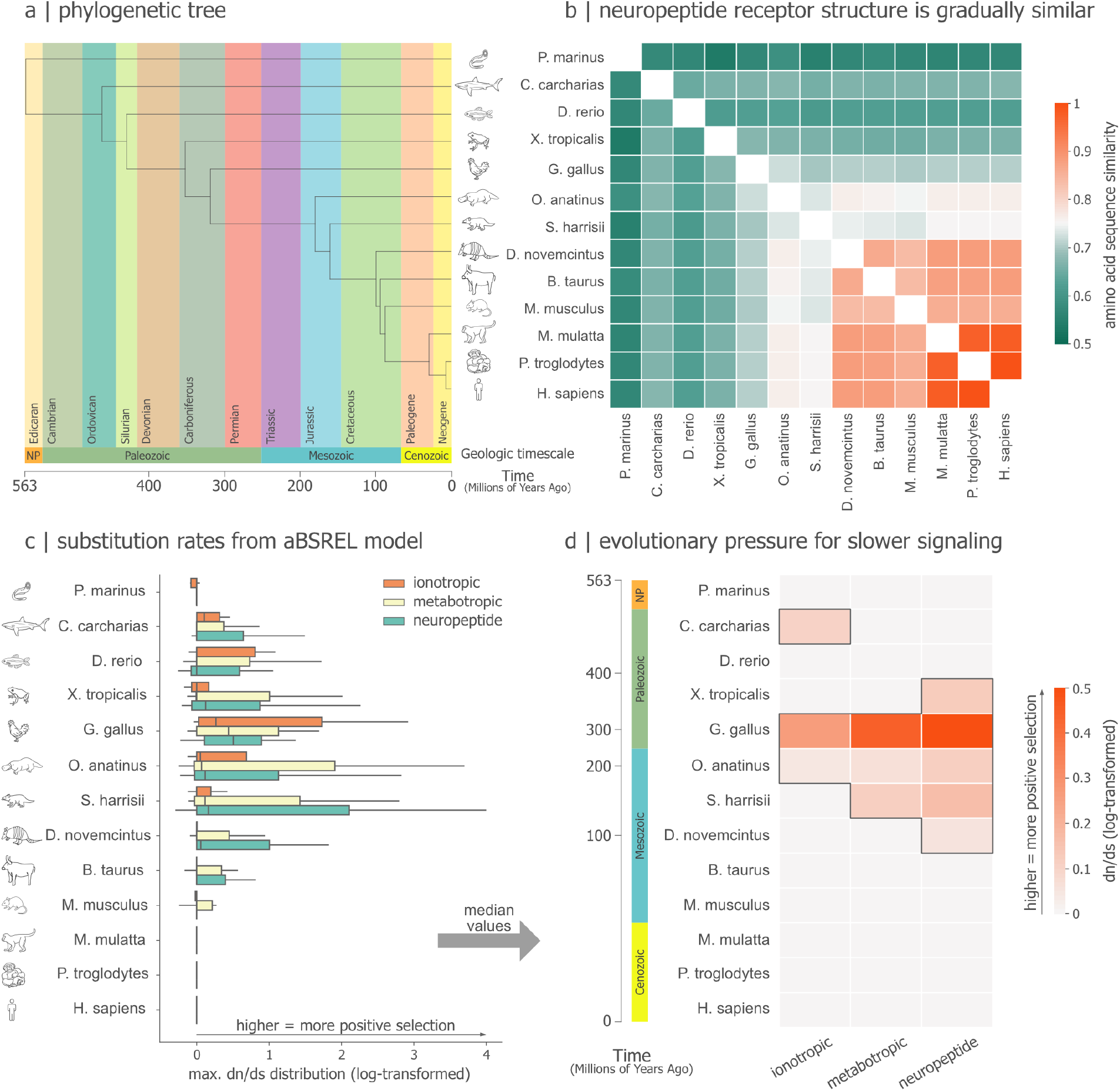
Neuropeptide signaling across phylogeny. (a) Phylogenetic lineage of human evolution for 13 species, each representative of a specific era in human evolution (e.g. first vertebrate, first tetrapod, etc.) [146]. (b) Amino acid sequence similarity of neuropeptide receptors across evolution. (c) Codon-based substitution rates for receptor genes, stratified by ionotropic and metabotropic neurotransmitters, as well as neuropeptides. Boxplot lines represent median and whiskers represent 25% and 75% quantiles. (d) Median substitution rates for different types of neuropeptide and neurotransmitter receptors. All values are logtransformed. Table S2 shows the exact statistical estimated and fitted parameters of the aBSREL model.

We investigate the evolutionary moments when neuropeptide receptors were preferentially selected during evolution, i.e., events of positive selection throughout our lineage. To contextualize neuropeptide signaling evolution, we additionally investigate convergences of other signaling receptors, namely ionotropic and metabotropic neurotransmitter receptors (see *Methods* for list of genes). We apply an adaptive branchsite random effects likelihood (aBSREL) model to parse the rate of non-synonymous versus synonymous changes (substitution rate dn/ds) in the genome as a measure of positive selection (see *Methods*) [156]. In other words, the aBSREL model allows us to know when tendencies to proliferate receptor genes existed in our evolution. Fig 6b shows the distribution of log-transformed substitution rates for ionotropic, metabotropic neurotransmitter, and neuropeptide receptor categories for our ancestor species. In this context, substitution rates above 0 indicate positive selection, values at 0 indicate neutral evolution, and negative values indicate purifying selection.

Fig. 6c shows the distribution of substitution rates among signaling types. We observe that the genetic evolution of signaling genes is characterized by periods of intense selection followed by relaxation at different rates for each signaling type. Statistical testing confirms this effect of signaling type on substitution rates (Friedman *χ*^2^ = 7.54, p = 0.023, Kendall’s W = 0.29), indicating moderate variability in positive selection between ionotropic, metabotropic, and neuropeptide genes. Collapsing the distributions onto their median (Fig. 6d), we observe that signaling genes altogether peak around G. gallus and O. anatinus. Once again, we observe that each signaling type differs in their dynamic, with positive selection for ionotropic genes slowing down before metabotropic neurotransmitter and neuropeptide genes. Taken together, this suggests a demand for signaling genes around the transition from amniotes (G. gallus) to mammals (O. anatinus), with sustained selection for slower signaling during early mammalian evolution. Interestingly, compared to neurotransmitter receptors, we find that peptide receptors evolve over a more protracted time scale. This evolutionary differentiation of genes is also mirrored by their anatomical localization: when inspecting neuropeptide genes that are significantly positively selected, we observe that genes selected in earlier evolutionary branches are now predominantly found in subcortex, e.g., APLNR, EDNRB, NPR2 genes. In contrast, neuropeptide genes that show selective pressure in more recent stages of evolution, such as MCHR1, VIPR2, NPFFR2, GRPR, are now found to be expressed foremost in the cortex. Altogether, these results suggests that the cortex-subcortex differentiation observed in the previous sections may have an evolutionary origin and underscore the concomitant emergence of neuropeptide signaling, and mammalian cortical development and higher cognitive function.

### Sensitivity and Specificity

As a final step, we seek to further establish the validity and robustness of the results. In addition to the correlations between gene expression and PET-estimated protein density shown in Fig. 3, we perform three other sanity checks based on domain knowledge about neuropeptides. First, we find that neuropeptides previously known to display sexual dimorphism – such as relaxin [34], melanin-concentrating hormone [145], and oxytocin [7, 137] – also have differential expression between male and female donors (Fig. S2), consistent with their role in reproduction [30, 78, 111]. Second, we find significant spatial correlations between endothelin receptors, known to regulate blood supply to stay within healthy physiological ranges [4, 91, 105, 117], and regional blood flow estimated using arterial spin labeling (ASL; Fig. S4). Third, using only Neurosynth maps related to “feeding” and “eating”, we find significant enrichment for neuropeptides known to modulate appetitive behaviour (Fig. S5d), such as leptin [44, 83, 89, 104] and calcitonin [87, 122, 165]. In addition to these specificity checks, analyses were contrasted with multiple null models, including spatial autocorrelation-preserving randomization and matched gene nulls (see *Methods*). Finally, we repeat the analyses using an alternative parcellation based on anatomical landmarks [25], finding an equivalent gene expression distribution (Fig. S3).

## DISCUSSION

Neuropeptides constitute a fundamental chemical communication mechanism throughout the nervous system and the entire body [22, 75, 159, 161]. Unlike neurotransmitters, neuropeptides are released under sustained neural activity and remain diffused through extracellular space for longer periods [22, 174, 183]. Their slow, global presence in circuits has drawn increasing attention to them as key regulators of states beyond local synaptic signaling [72, 74, 97, 115, 174]. Despite their importance, neuropeptide signaling remains uncharted territory [161], with only recent efforts trying to map its presence at a system level [134, 157]. Here, we comprehensively mapped whole-brain human neuropeptide signaling across multiple levels of description; from molecular and cellular embedding to mesoscale connectivity and macroscale cognitive specialization. As we elaborate in this section, neuropeptide signaling is highly organized and leaves an indelible mark on several features of brain structure and function.

Spatially, neuropeptide systems exhibit a striking anatomical organization across cortical and subcortical regions. For instance, receptors for somatostatin, vasoactive intestinal peptide (VIP), and melanin-concentrating hormone (MCH) are predominantly expressed in the cortex, while other families, such as neuropeptide Y (NPY), opioid, natriuretic, and endothelin receptors, are expressed across both cortical and subcortical regions. In contrast, receptors for neuropeptides like oxytocin and calcitonin are more restrained to subcortical nuclei. This anatomical gradient suggests a potential specialization in the roles played by different neuropeptides, with some potentially modulating cortical and subcortical circuits together, while others act more focally within one structure. As we outline below, the confluence of gross anatomical localization, signaling environment and prolonged time scale of effect may altogether confer the capacity to modulate a variety of bodily and cognitive functions.

At the cellular level, we show that neuropeptide receptors tend to co-localize with predominantly metabotropic, and, to a lesser extent, ionotropic neurotransmitter systems. This arrangement is consistent with the notion that neuropeptides primarily act as a global modulation architecture to facilitate or restrict local G protein-coupled plasticity from metabotropic neurotransmitter activity [120, 141]. This is akin to previous theoretical frameworks of meta-plasticity and three-factor learning, where a third modulatory factor influences traditional Hebbian learning [1, 50, 120]. To a lesser degree, neuropeptides may also prime local ionotropic activity, enabling or disabling a milieu of excitable cells to promote spike timing dependent plasticity [120]. Here, previous literature has elucidated the computational flexibility that arises from co-transmission of fast (ionotropoic) and slow (neuropeptide) signaling [116, 163], providing a biological grounding of the multiplexed signaling environment observed in the brain. Together, the manifold of communication protocols may thus endow a repertoire of functions, ranging from fast ionotropic communication to slow, sustained neuropeptide state signaling to allow for both immediate responses and long-term adaptability [19, 42, 120].

At the mesoscale, we find pervasive involvement of neuropeptide signaling in whole-brain circuits. We find that the alignment between an area’s neuropeptide ligand profile and the receptor profile of its connected neighbours shapes their functional connectivity. Indeed, several recent reports suggest that structure-function coupling in the brain is regionally heterogeneous and can be optimized by introducing local biological annotations into connectome models [12, 67, 94, 177]. The present findings are a salient example of how neuropeptide signaling can support a wide spectrum of communication mechanisms. In particular, we find that certain peptide families, like natriuretic peptides, fit neurovascular haemodynamic connectivity better, others like somatostatin fit electrophysiological oscillatory connectivity better, and others, like endothelin and opioids, fit both. Many of these associations are supported by previous research; for instance, multiple studies have reported evidence of opioid modulation of both electrical and neurovascular activity [28, 32, 58, 61]. More broadly, linking neuropeptides and connectivity highlights how information about neuropeptide receptors and ligands can be used to extend the scope of inquiry from local to global communication. These findings are part of an emerging literature showing that the configuration of synaptic connections between neurons is closely related to their molecular and cellular features [8, 13, 17, 56, 68, 148, 158].

At the systems level, we show that neuropeptides colocalize with regional patterns of cognitive specialization. Specifically, we find that subcortical neuropeptide receptors are associated with behavioral states related to homeostasis and state regulation, including reward anticipation, reinforcement learning, stress, anxiety, sleep, and eating. In contrast, cortical neuropeptide receptors are linked to functions related to cognition, such as visuospatial attention, multisensory integration, and intention. This underlines the role of neuropeptides as systemic regulators, and is one of several recent studies investigating how molecular features shape regional cognitive specialization [49, 66, 67, 185]. The concomitant molecular-anatomical differentiation of function, with cortical peptides supporting cognitive and subcortical modulating sensory-cognitive processes, may potentially reflect the evolutionary history of the nervous system, a question we consider next.

From an evolutionary perspective, refinement of neuropeptide and metabotropic neurotransmitter signaling appears to increase during the transition from amniotes to early mammals. We find heightened selection for neuropeptide receptor genes during this period, with those positively selected before the emergence of the cortex predominantly localized in subcortex. With the advent of mammals, later-selected neuropeptide receptors are now found in cortex, supporting the idea that that neuropeptide signaling location can be in part explained by the timing of its evolutionary pressure. Moreover, we find that the evolutionary pressure for metabotropic and neuropeptide signaling is more protracted around the advent of mammals than for ionotropic, suggesting a continued refinement of their function during this time period. Indeed, neurons might have opted for neuropeptides due to their ability to produce multiple peptides from one single precursor gene, thereby facilitating signaling diversity while being able to regulate peptide activity through enzymatic modifications that respond to cellular conditions [22, 183]. Together with the knowledge that peptides are an ancient signaling mechanism [134, 146], we hypothesize that neuropeptides have been co-opted for a wide range of functions throughout evolution, ranging from basal cellular regulation to higher cognitive computations.

While our study offers promising insights, several methodological limitations should be acknowledged. First, our analyses use bulk microarray gene expression data from the Allen Human Brain Atlas (AHBA), which, despite being the most spatially comprehensive dataset currently available, is limited in resolution and may not perfectly correspond to protein density [65, 95]. We attempted to address this limitation using a rigorous quality control selection and cross-reference selected genes with RNA-seq data from the Human Protein Atlas S1. In addition, we found excellent correspondence between gene expression and protein density for two opioid receptors for which PET measurements were available. Looking ahead, emerging technologies such as GPCR-activation–based sensors that track peptide signaling across the whole brain [180], and advances in PET molecular imaging with refined radiochemistry [140], may offer more precise measurements in the future. Second, gene expression is known to vary with demographic factors such as sex, age, and environmental influences [33, 179]. We observed sex-based differences in neuropeptide receptor expression, underscoring the need for future research into individual-level variability. As more comprehensive datasets become available, particularly those capturing more diverse populations, these factors will open up exciting new avenues for research in the future. Third, our focus on gene transcription primarily captures postsynaptic and local presynaptic signaling. To assess potential long-range neuropeptide signaling, we analyzed the overlap of ligand and receptor expression between regions, estimating their roles in connectivity. While this approach recovered several previously reported findings, certain relationships, such as NPY-NPY1R and BOLD signaling [150, 171], remained undetected, warranting cautious interpretation. Fourth, the structural connectome used in our analysis was reconstructed from diffusion-weighted imaging, which is prone to false positives and negatives [47, 96]. Although we minimized this limitation by employing modern multi-shell diffusion data and state-of-the-art tractography, the inherent challenges of tractography remain. Future efforts will need to address these challenges, and we make our code and data openly available to facilitate further research and validation of our results.

In conclusion, we show that neuropeptides represent an omnipresent component of signaling within the human brain, acting at multiple scales to influence both local synaptic dynamics and global brain states. Their spatial distribution across cortical and subcortical regions reflects their diverse roles from modulating cognition to bodily states. The evolutionary preference for slower neuropeptide signaling, alongside metabotropic neurotransmitters, highlights its fundamental role in the development of the mammalian brain, putatively contributing to the complexity and adaptability of modern neural circuits. Neuropeptides, through their long-lasting and widespread effects, can thus be seen as key players in the coordination of both immediate and enduring neural and behavioral responses.

## METHODS

The code to perform all analyses is available at https://github.com/netneurolab/ceballos_neuropeptide-signaling, with data available at https://osf.io/4rsz9/.

### Gene expression data

Gene expression data was acquired from the Allen Human Brain Atlas (n=6, 1 female, ages 24-57, three of Caucasian ethnicity, two African American, one Hispanic, 71) using the *abagen* toolbox (v0.1.4, https://github.com/rmarkello/abagen, 98). Detailed information about data acquisition can be found in [71]. Preprocessing of gene expression data closely followed the procedures described in [3]. Microarray probes for all individuals were fetched from https://human.brain-map.org/ and re-annotated to match the gene symbol ID and name from the latest version of NCBI [46], found under https://ftp.ncbi.nih.gov/gene/DATA/GENE_INFO/Mammalia/. Probes not matched to a valid ID were discarded. The remaining microarray probes were subject to intensity based filtering, where probes with intensity less than background noise in less than 20% of samples across individuals were excluded. For genes that had multiple probes assigned, the probe with the highest average Pearson correlation to the RNA-seq data from two individuals in the dataset was selected. This ensured biological plausibility of the micro-array measurements wherever possible [106]. Tissue sampling locations from the original T1w scans of individuals were then non-linearly registered to MNI 152 1mm space as in https://github.com/chrisgorgo/alleninf. Probe samples were assigned to brain regions in a standard atlas consisting of the seven-network Schaefer 400 atlas [147], the Melbourne Subcortex Atlas [167], and the CIT168 atlas [118] for the hypothalamus, thresholded at 0.5 probability. Samples were then assgined to a brain region if their MNI coordinates were within 2 mm vicinity. To reduce the potential of misassignment, sample-to-region matching was constrained by hemisphere and gross structural divisions (i.e., cortex, subcortex/brainstem, and cerebellum, such that e.g., a sample in the left cortex could only be assigned to an atlas voxel in the ipsilateral side). If a brain region was not assigned a sample based on that procedure, every voxel in the region was mapped to the nearest tissue sample from the individual in order to generate a dense, interpolated expression map. The mean of these expression values was taken across all voxels in the region, weighted by the distance between each voxel and the sample mapped to it, in order to obtain an estimate of the parcellated expression values for the missing region. Inter-subject variation was addressed by normalizing sample expression values across genes using a robust sigmoid function as in [53]. Normalized expression values were then rescaled from 0 to 1 by min-max normalization. Gene expression values were subsequently normalized across regions using an identical procedure. All available tissue samples were used in the normalization process regardless of whether they were assigned to a brain region. Tissue samples not matched to a brain region were discarded after normalization. Samples assigned to the same brain region were averaged separately for each individual, which resulted in six expression matrices, one for each individual, with rows corresponding to brain regions and columns corresponding to genes. All matrices were finally mean-averaged across individuals, resulting in a single matrix representative of the expression level of a particular gene in a given region.

### Receptor PET data

PET-derived receptor density maps were collated by Hansen et al. [67] and downloaded from neuromaps (https://github.com/netneurolab/neuromaps [99]) for 16 neurotransmitter receptors and transporters across 8 neurotransmitter systems. These include dopamine (D2 [144]), serotonin (5-HT1A [14], 5-HT1B [54], 5-HT2A [14], 5-HT4 [14], 5-HT6 [130, 131]), acetylcholine (*α*4*β*2 [73], M1 [108]), glutamate (mGluR5 [38]), GABA (GABA_*A*_ [113]), histamine (H3 [55]), cannabinoid (CB1 [114]), and opioid (MOR [169], KOR [178]). Methodological details about each tracer can be found in the neuromaps documentation (https://netneurolab.github.io/neuromaps/listofmaps.html) or in the original report of the receptor library [67]. Volumetric PET images were parcellated according to the Schaefer 400-region [147], Melbourne Subcortex S4 [167] atlas as well as the hypothalamus delineation from the CIT168 atlas [118].

### Dominance analysis

To associate neurotransmitter maps with neuropeptide receptors, we opted to use dominance analysis [6], a multilinear regression technique, for two reasons. First, it allows us include every available neurotransmitter receptor map at once to predict the spatial expression of neuropeptide receptors. Consequently, we are able to compensate the heterogeneous affinity of PET tracers across cortical and subcortical structures – see e.g. discussion about [11C]raclopride and [11C]flb457 usage for D2 localization in cortex and subcortex [40, 45, 80]. Second, dominance analysis quantifies the prediction performance of a neurotransmitter receptor by building subsets of the full multiple linear regression model and calculating the average contribution of a neurotransmitter receptor when being included into all possible models that include it. This results in a measure of the total (general) dominance [6] a neurotransmitter receptor has on the spatial expression of a neuropeptide receptors. The sum of all total dominances is equivalent to the *R*^2^ of the full model, which means that the total dominance of each neurotransmitter receptor can be normalized by its sum to obtain the percentage it contributes to the total *R*^2^, also known as relative contribution can then be turned into a percentage of the total spatial variance explained by a neurotransmitter receptor. In other words, we are able to answer the question of how much more in line are the distributions of neurotransmitter and neuropeptide receptors when including a particular neurotransmitter receptor into the multilinear model. Finally, to test whether there is a difference in the percentage explained by ionotropic and metabotropic neurotransmitter receptor densities, we first averaged the total dominance of each and then compared the relative contribution of the two classes to the total sum *R*^2^.

### Connectomic data

The map of hypothalamic connectivity was derived from a group-averaged functional connectivity template, constructed using high-resolution 7-Tesla resting-state fMRI data from 20 unrelated healthy participants [27, 64, 154]. Cortical regions were defined by the Schaefer 400 atlas [147]. Hypothalamic delineation was defined using the probabilistic CIT168 subcortical nuclei atlas, thresholded at a probability > 0.5 [118]. Functional connectivity (FC) between the hypothalamus and cortex was derived from Pearson correlations between the time series of these regions. Specific preprocessing protocols for imaging of deep nuclei, including physiological noise reduction, were used to enhance the signal quality [64]. For further details on the construction of this connectome, we refer to the original publications [27, 154].

The template fMRI and dMRI connectomes used in our structure-function coupling analysis were constructed from a cohort of 1065 participants from the Human Connectome Project (HCP) S1200 dataset using a group consensus method [15, 16]. Code for the consensus method is found in the netneurotools package (https://github.com/netneurolab/netneurotools/).

Resting-state fMRI data were acquired using a multiband echo-planar imaging (EPI) sequence with the following parameters: TR = 720 ms, TE = 33.1 ms, flip angle = 52°, and voxel size = 2 mm isotropic. Each participant underwent two 15-minute fMRI sessions, one in each phase encoding direction (left-right and right-left). Detailed acquisition parameters and scanning protocols can be found in the original HCP papers [43, 175]. The functional imaging data were preprocessed using the HCP Minimal Preprocessing Pipeline [59], which includes motion correction, intensity normalization, distortion correction, and registration to the standard MNI space. Surface-based registration to a standard cortical surface was also applied to facilitate cross-subject comparisons.

Diffusion-weighted imaging (dMRI) data were acquired using a spin-echo EPI sequence with the following parameters: TR = 5520 ms, TE = 89.5 ms, flip angle = 78°, voxel size = 1.25 mm isotropic, and bvalues of 1 000, 2 000, and 3 000 s/mm^2^. Three sets of diffusion data with distinct gradient directions (90 in total) were collected. Detailed acquisition parameters can also be found in the original HCP papers [43, 175]. Briefly, the diffusion data were preprocessed using HCP Minimal Preprocessing Pipelines [59], which includes intensity normalization, distortion correction (“topup”), eddy current and subject head motion correction (“eddy”), and gradient nonlinearity correction. We downloaded bedpostX-derived fiber orientation density function (fODF) estimation from the HCP S1200 release, and ran probabilistic tractography using probtrackx2 to estimate the structural connectivity between brain regions (dwi_probtrackx_dense_gpu command in Qunex v0.98.0; 79).

Resting state MEG data was acquired for a subset of *n* = 33 participants (age range 22-35 years). The data consists of resting state scans of approximately 6 minutes long (sampling rate = 2,034.54 Hz; anti-aliasing filter low-pass filter at 400 Hz) and noise recordings for all participants. Preprocessing was conducted with Brainstorm software (version 220420; 164). The pipeline includes applying notch filters (60, 120, 180, 240, 300 Hz), a high-pass filter (0.3 Hz), bad channel removal, and automatic artifact removal (including heartbeats, eye blinks, saccades, muscle movements and noisy segments). Heartbeats and blinks were detected using electrocardiogram (ECG) and electrooculogram (EOG), respectively. Saccade and muscle activity were detected as low-frequency (1 to 7 Hz) and high-frequency (40 to 240 Hz) components, respectively. Detected artifacts were removed using Signal-Space Projections (SSPs; 172). Preprocessed data at the sensor level underwent a source estimation procedure using a linearly constrained minimum variance (LCMV; 176) beamformer on HCP fsLR4K surface. Surface-level data was then parcellated using the Schaefer 400 atlas [147]. Finally, MEG functional connectivity was derived from amplitude envelope correlation (AEC; 20) for six canonical electrophysiological bands: *δ* (2-4 Hz), *α* (8-14 Hz), *β* (15-29 Hz), low *γ* (30-59 Hz) and high *γ* (60-90 Hz).

### Structure-function coupling

Structure-function coupling measures the overlap between functional connectivity and brain communication, i.e., the propensity for brain regions to communicate with each other via anatomical connections [94, 160]. To evaluate communication at multiple levels, we employ a range of protocols previously used to predict functional connectivity [94, 177]. These include shortest path length, navigation efficiency, search information and communicability, which are thought to reside on a spectrum from decentralized, diffusion-like (e.g., communicability) to centralized, routing-like (e.g., shortest path routing) communication processes [5, 11, 149]. We determined a baseline fit between structural and functional connectivity by using the above communication matrices (non-diagonal upper triangular elements) as linear predictors for each of the seven functional connectivity matrices (one haemodynamic and six electrophysiological). The global structure-function coupling is then interpreted as the adjusted goodness-of-fit *R*^2^ of the overall model [60, 177, 187], thereby accounting for the multiple predictors. We repeat this procedure, now re-weighing the otherwise binary structural connectome matrix with the outer product of neuropeptide receptor and their cognate ligand gene expression. In other words, for two brain regions A and B, we annotate the corresponding edge that connects A and B as the product of receptor and ligand gene expression, thereby delineating the the concordance between receptor and ligand availability between two connected regions. Then, using the annotated structural connectome, we again derive structural communication predictors that we fit to all functional connectivity modes, resulting in new structure-function coupling values that we can benchmark to the unannotated fits.

### Neurosynth meta-analytic term data

Term-specific associations between voxels and cognitive processes were obtained from data in Neurosynth [186], a meta-analytic tool that synthesizes results from more than 14,000 published functional MRI studies by searching for high-frequency keywords (such as “pain” and “attention”) that are published alongside functional MRI voxel coordinates. We model activation based on activation likelihood estimation (ALE; 41), which estimates the consistency in which a voxel is reported for a given term across studies, highlighting brain regions with increased likelihood to be associated with that term. Although more than 10,000 functional processes are reported in Neurosynth, we focused primarily on cognitive function and, therefore, limit the terms of interest to cognitive and behavioral terms. These terms were therefore cross-referenced with the Cognitive Atlas, a public ontology of cognitive science [126], which includes a comprehensive list of neurocognitive processes. We used 125 terms, ranging from umbrella terms (“attention” and “emotion”) to specific cognitive processes (“visual attention” and “episodic memory”), behaviors (“eating” and “sleep”) and emotional states (“fear” and “anxiety”). The coordinates reported by Neurosynth were parcellated according to the Schaefer 400-region [147] and Melbourne Subcortex S4 [167] atlases. Additionally, we included the hypothalamus delineation from the CIT168 atlas [118]. The full list of all terms is shown in Supplementary Fig. S5.

### Random matched gene null set

To assess whether the observed spatial autocorrelation patterns of neuropeptide receptor genes were unique or consistent with random gene expression patterns, we instantiated a random sampling approach to contrast our results to other genes that match the spatial expression of a given neuropeptide receptor based on two criteria: spatial autocorrelation and expression value distribution. This procedure was repeated 10 000 times for each of the 38 neuropeptide receptors that passed our quality control protocol. We outline our approach below.

First, we randomly selected *n* = 100 genes from all genes identified in the Allen Human Brain Atlas to create a random subset of genes. This set excluded neuropeptide receptor and precursor genes. To ensure a close match in spatial autocorrelation, the Moran’s I of each gene in the subset was computed and compared to the corresponding value of the neuropeptide receptor gene [13]. The Kolmogorov-Smirnov (K-S) statistic was then estimated to assess how well the expression distribution of the random genes matched that of the neuropeptide receptors. For each neuropeptide receptor gene, we ranked the difference between its Moran’s I and that of each null gene, as well as the K-S statistic between their expression distributions. The null gene with the best average rank between the two criteria was then chosen as closest match and recruited into the null distribution.

### Spatial autocorrelation-preserving null

For all analysis comparing the spatial distribution between brain maps, we employed BrainSMASH [23, 24, 100] as implemented in the neuromaps [99] library. Briefly, BrainSMASH is a null model to generate surrogate brain maps that match a target spatial autocorrelation [24]. The model entails two steps: 1) random permutation of values within a designated brain map and 2) smoothing and re-scaling to restore the spatial autocorrelation characteristic of the target data. In the first step, values within the brain map are randomly permuted. The permuted data is then transformed to reintroduce spatial autocorrelation. This transformation involves combining the permuted data and a normally distributed random variable. Then, ratios in this combination are optimized using least-squares to match the spatial variance patterns (variograms) of the target and surrogate data.

### Partial Least Squares Correlation

We applied Partial Least Squares Correlation (PLSC) to investigate the relationship between behavioral term maps derived from Neurosynth (*X* matrix) and the gene expression profiles of neuropeptide receptors (*Y* matrix) [102, 103]. Both matrices were centered and scaled to unit variance in each column to ensure comparability. PLSC identifies latent variables that capture the maximum shared variance between the two datasets by performing singular value decomposition (SVD) on the cross-covariance matrix *X*^*T*^ *Y* . This process was carried out using a Python implementation of PLSC (*behavioral_pls*), openly available at https://github.com/netneurolab/pypyls.

The decomposition yields singular values that reflect the strength of the covariance explained by each latent variable pair, and singular vectors that weigh the contribution of the original variables (i.e., neuropeptide receptor expression and term activations). The latent weights were applied to the respective matrices to obtain the PLSC scores, which represent the data projected onto the latent components. Loadings, representing the contribution of each map to the latent variable, were computed by assessing the similarity of each map to its respective score using Pearson correlation. This allows us to interpret how individual behavioral term maps and gene expression profiles contribute to each latent component.

To assess the significance of the identified latent components, we employed two distinct permutation testing strategies. First, we generated a set of random null maps that preserved the spatial autocorrelation properties of the neuropeptide receptor maps and shuffled the *Y* matrix accordingly [66]. Second, we exchanged the *Y* matrix with a set of random matching genes that maintained key properties of the gene expression data (see *Random matched gene null set*). For each permutation test, we computed the singular values of the cross-covariance matrix for the shuffled data, thereby generating null distributions of singular values. The significance of the actual singular values was determined by comparing them to the null distributions derived from both procedures.

To ensure the generalizability of the PLSC results, we performed cross-validation by randomly splitting the observations into training and test sets 1 000 times [107, 132]. The training set comprised 70% of the observations, and the PLSC was repeated using this subset. The goodness-of-fit was evaluated by calculating the correlation between the scores of *X*_training_ and *Y*_training_. The projection weights derived from the training set were then applied to the remaining 30% in the test set, transforming the test observations into scores. We similarly assessed the correlation between the scores of *X*_test_ and *Y*_test_ to evaluate how well the PLSC model generalized to unseen data.

### Evolutionary analysis

To investigate signatures of positive selection in ionotropic, metabotropic, and peptide signaling pathway genes, we used a modified procedure based on the original analysis of [146]. We focused on receptor genes associated with ionotropic (‘CHRND’, ‘GRIA1’, ‘CHRNA1’, ‘HTR3A’, ‘GRIK1’, ‘CHRNG’, ‘GABRG1’, ‘GABRA1’, ‘CHRNE’, ‘GRIN1’, ‘GABRA2’, ‘GABRG2’, ‘CHRNB1’), metabotropic signaling (‘GABBR1’, ‘HRH2’, ‘DRD2’, ‘ADRB2’, ‘CHRM2’, ‘DRD4’, ‘HRH4’, ‘ADRA1A’, ‘CHRM1’, ‘ADRB1’, ‘MTNR1A’, ‘HTR4’, ‘HTR2A’, ‘HRH1’, ‘DRD1’, ‘GABBR2’, ‘GRM1’, ‘ADRB3’, ‘CHRM3’, ‘ADRA2A’, ‘HTR1A’, ‘DRD3’, ‘HRH3’), and compared those to the ensemble of neuropeptide receptor genes previously used in order to contextualize our results.

Orthologous amino acid sequences (AAS) and coding sequences (CDS) for the selected genes were manually downloaded from the NCBI database for thirteen vertebrate species: Homo sapiens (human), Pan troglodytes (chimpanzee), Macaca mulatta (rhesus macaque), Mus musculus (house mouse), Bos taurus (cattle), Dasypus novemcinctus (nine-banded armadillo), Sarcophilus harrisii (Tasmanian devil), Ornithorhynchus anatinus (platypus), Gallus gallus (chicken), Xenopus tropicalis (western clawed frog), Danio rerio (zebrafish), Carcharodon carcharias (great white shark), and Petromyzon marinus (sea lamprey). These species represent key evolutionary lineages within vertebrates and provide a comprehensive framework for identifying evolutionary signatures across a broad phylogenetic spectrum [146].

For each gene, the collected AAS and CDS from the different species were compiled into a single protein FASTA file and a CDS transcript FASTA file. The protein sequences in each file were aligned using MUSCLE (v5.1; 39) to create multiple sequence alignments for each receptor gene. These alignments were then translated into codon-based alignments using PAL2NAL (v14; 162), which converts protein sequence alignments into nucleotide alignments while preserving codon structure. The resulting codon-alignment files were further processed by inserting dummy sequences for missing orthologs, renaming sequence identifiers to abbreviated taxon names (e.g., ‘Homo sapiens’ became ‘hsapiens’), and converting the FASTA format codon alignments to PHYLIP sequential format files using TriFusion (1.0.1; 153). We report the similarity of amino acids from the PHYLIP sequential format files as computed in TriFusion.

In addition, a vertebrate species tree was generated using TimeTree [88], incorporating data from published studies to produce a reliable species tree. The tree was downloaded in Newick format, with branch lengths manually removed to match the format required for subsequent analyses. Taxon names in the tree file were shortened to match those in the codon-based alignments.

Positive selection analyses were conducted using the adaptive branch-site random effects likelihood (aBSREL) model from the HyPhy package (v2.5.52; 86, 127). The aBSREL method is an exploratory tool for detecting episodic positive selection across all branches in an evolutionary tree without the need for specifying foreground branches a priori. This model compares a full, alternative model against a null model and performs a Likelihood Ratio Test (LRT) to identify the magnitude that the empirical results exceed chance (see [156] for a detailed description of the aBSREL method). Our main results are based on the categorical median substitution rates – also known as dn/ds or omega ratios – between ionotropic, metabotropic, and peptide receptor genes. Substitution rates are informative in this context as they provide insight into the evolutionary pressure exerted on genes, allowing us to differentiate between genes that have undergone positive selection and those that have remained under purifying selection at a particular branch. By comparing these rates across signaling types, we can infer the relative evolutionary dynamics and selective pressures that have shaped neuropeptide functionality.

### Friedman test

To investigate whether the substitution rates (dn/ds) varied significantly across the three signaling categories – ionotropic, metabotropic, and peptides – a non-parametric Friedman test was performed [51]. This test was chosen due to its suitability for repeated measures data, where dn/ds values were modeled for 13 branches representing species in the human lineage. We chose to model the categorical difference between signaling types, treating each branch as a repeated measure across categories, to account for phylogenetic relatedness and shared evolutionary history. This approach enabled a robust assessment of differences in median substitution rates across categories without assuming normality in the data distribution. Significance was determined at p < 0.05, and effect size was calculated using Kendall’s W to quantify the strength of association between gene category and substitution rate variation [81].

### Human Protein Atlas gene expression

Human Protein Atlas gene expression data are based on RNA sequencing (RNA-seq) analysis of micropunch samples collected from the Human Brain Tissue Bank at Semmelweis University, Hungary, covering 190 distinct brain regions, areas, and subfields [155, 170]. RNA extraction was performed using the RNeasy Plus Mini Kit (Qiagen), followed by mRNA enrichment using ribosomal RNA depletion. RNA integrity was verified with the Experion RNA HighSens Analysis kit (Bio-Rad), with samples meeting a minimum RNA Integrity Number (RIN) and 260/280 absorbance ratio for inclusion. Sequencing libraries were prepared using Illumina TruSeq Stranded mRNA reagents and sequenced on the NovaSeq 6000 platform (paired-end, 150 bp). The reads were mapped to the human reference genome GRCh37/hg19 using Ensembl gene models (v92) and quantified with Kallisto (v0.43.1) [155, 189, 192]. After quality control, transcript expression levels were calculated as Transcripts Per Million (TPM), with further normalization performed using trimmed mean of M-values (TMM) with NOISeq and batch effect correction applied using limma [135]. The resulting normalized TPM (nTPM) were log-transformed (log-10) and mapped to the Destrieux Atlas [36] based on region name matches.

### Cerebral blood flow data

Cerebral blood flow data comes from the HCP-Aging dataset [69], estimated using Arterial Spin Labeling (ASL) [37, 190]. The acquisition is based on a pseudocontinuous arterial spin labeling (pCASL) and 2D multiband echo-planar imaging sequence [152]. pCASL data were acquired with labeling duration = 1500 ms and five post-labeling delays = 200 ms, 700 ms, 1200 ms, 1700 ms, and 2200 ms, containing 6, 6, 6, 10, 15 control-label image pairs, respectively. Other parameters related to this sequence include: spatial resolution = 2.5 x 2.5 x 2.5 mm^3^, TR/TE = 3580/18.7 ms. Two M0 images for blood perfusion quantification were acquired at the end of all the acquisitions. Preprocessing was done according to the HCP pipeline for ASL data (https://github.com/physimals/hcp-asl; 82). The resulting cerebral blood flow maps from *n* = 678 individuals were then averaged into a group template and mapped to the Schaefer 400 cortical [147] and the Melbourne Subcortex S4 [167] atlas.

## Acknowledgments

We thank Andrea I. Luppi, Vincent Bazinet, Moohebat Pourmajidian and Yigu Zhou for their comments and suggestions on the manuscript. EGC acknowledges support from the Molson Foundation and the Fonds de Recherche du Québec - Nature et Technologies (FRQNT). BM acknowledges support from the Natural Sciences and Engineering Research Council of Canada (NSERC), Canadian Institutes of Health Research (CIHR), Brain Canada Foundation Future Leaders Fund, the Canada Research Chairs Program, the Michael J. Fox Foundation, and the Healthy Brains for Healthy Lives initiative. The funders had no role in study design, data collection and analysis, decision to publish or preparation of the manuscript.

---

**TABLE S1.**
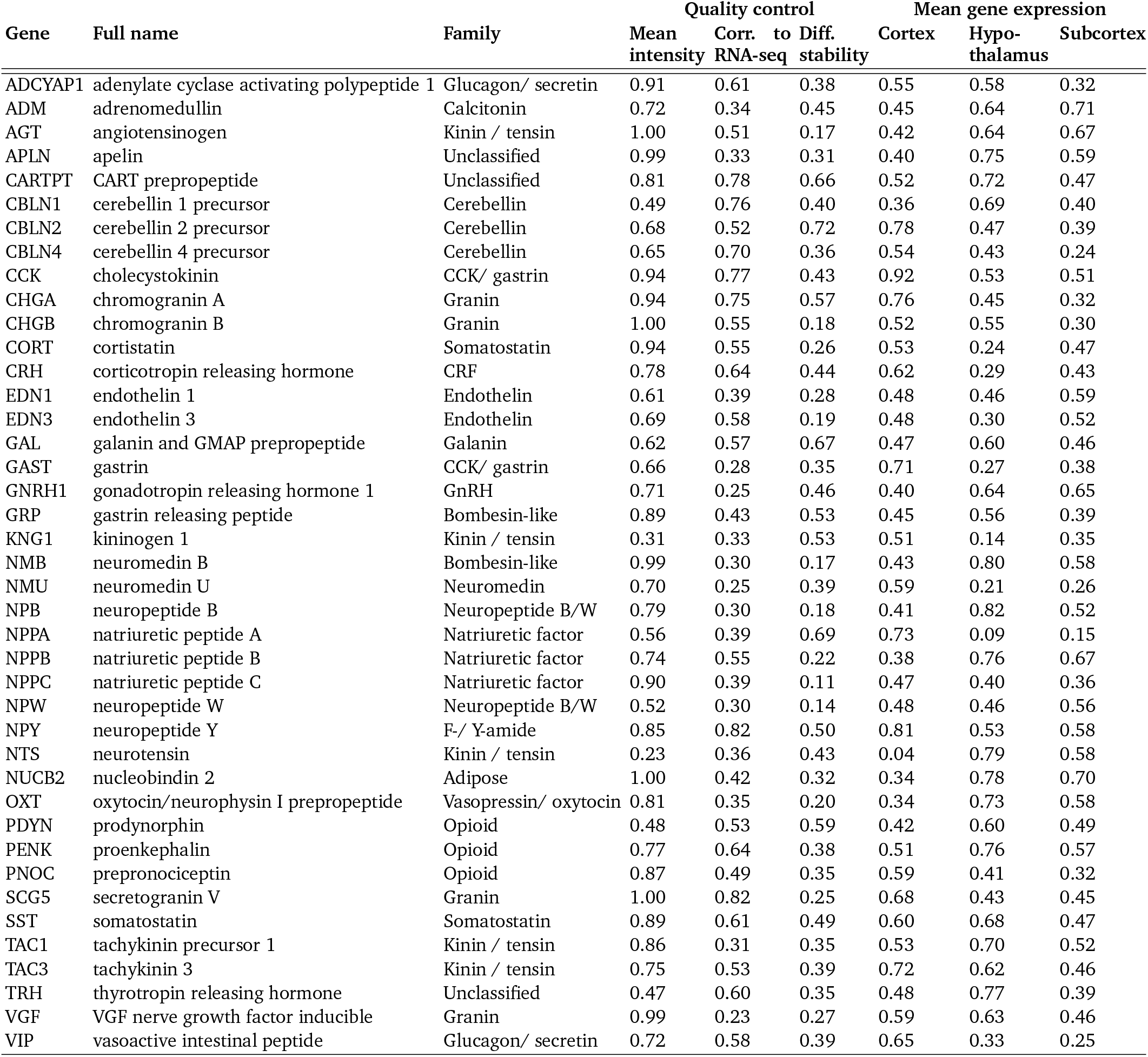
Neuropeptide precursor overview. Neuropeptide precursor distributions were estimated using gene expression from the Allen Human Brain Atlas [70, 71]. Just like in Table 1, gene transcripts were filtered based on multiple quality control criteria, including (1) mean intensity > 0.2, (2) correlation with RNA-seq > 0.2, and (3) differential stability > 0.1. For reference, mean expression within cortex, hypothalamus and subcortex are shown.

**TABLE S2.**
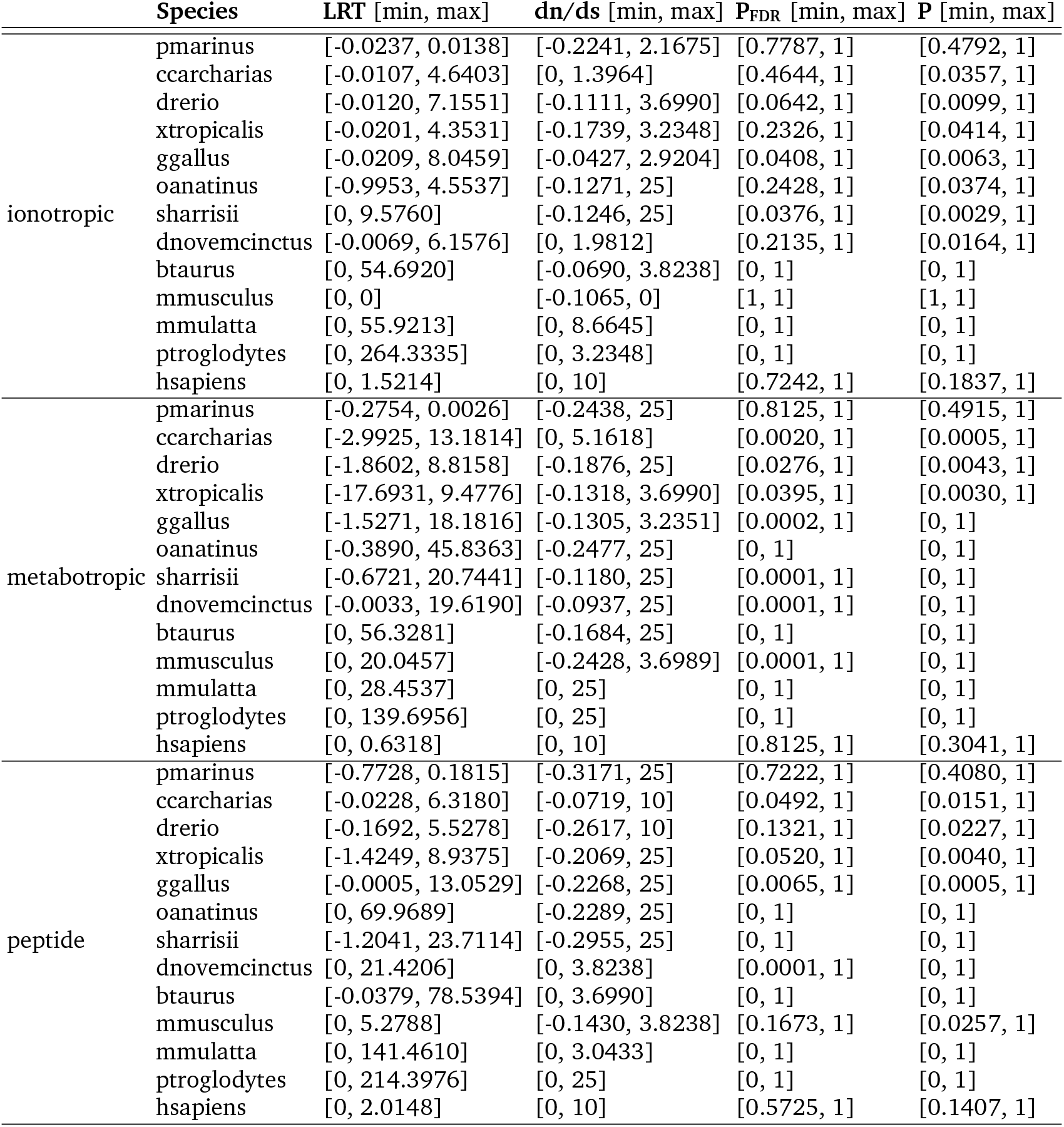
aBSREL results overview. Exact values of all ranges of likelihood ratio tests (LRT) and substitution rates (dn/ds) for each branch of the aBSRSEL model, stratified according to receptor type. These data are displayed as boxplots in Fig 6.

**Figure S1.**
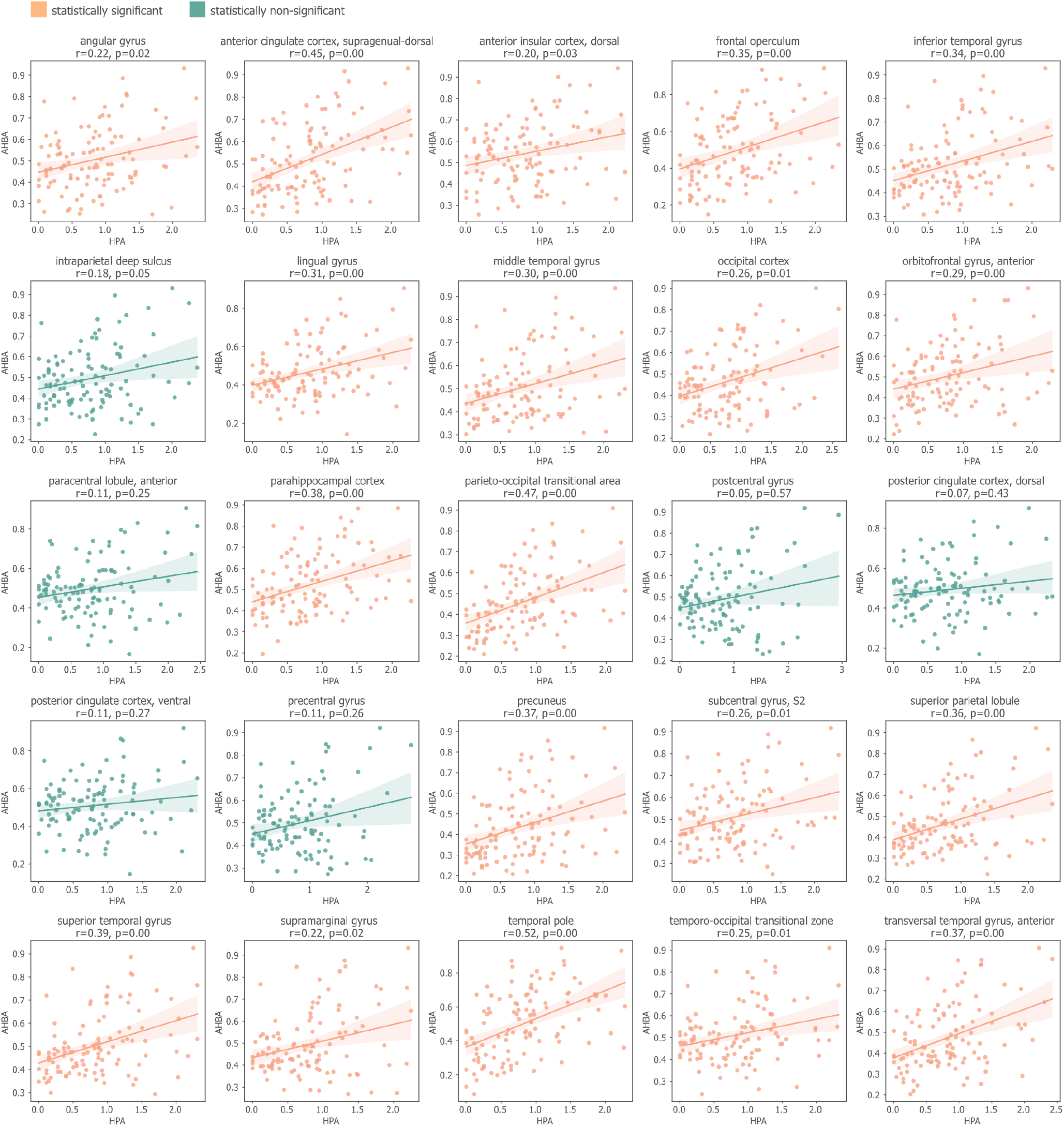
Comparisons with RNA-seq data from the Human Protein Atlas (HPA) To assess the consistency of spatial patterns of gene expression in the discovery data set (microarray bulk sequencing from the Allen Human Brain Atlas) [71], we systematically compare it with RNA-seq data from the Human Protein Atlas (HPA) [155, 170]. The AHBA data were parcellated according to the Destrieux atlas [36], and matched to the samples in HPA [155, 170], resulting in 25 regions of interest. Each panel corresponds to an anatomical region, with expression of neuropeptide receptor genes shown as points. The x-axis shows gene expression in HPA and the y-axis shows gene expression in AHBA. Orange and green plots indicate regions where linear associations are statistically significant and non-significant, respectively. We find positive correspondences between the datasets in all matched regions, and statistically significant correspondence in 19/25 regions.

**Figure S2.**
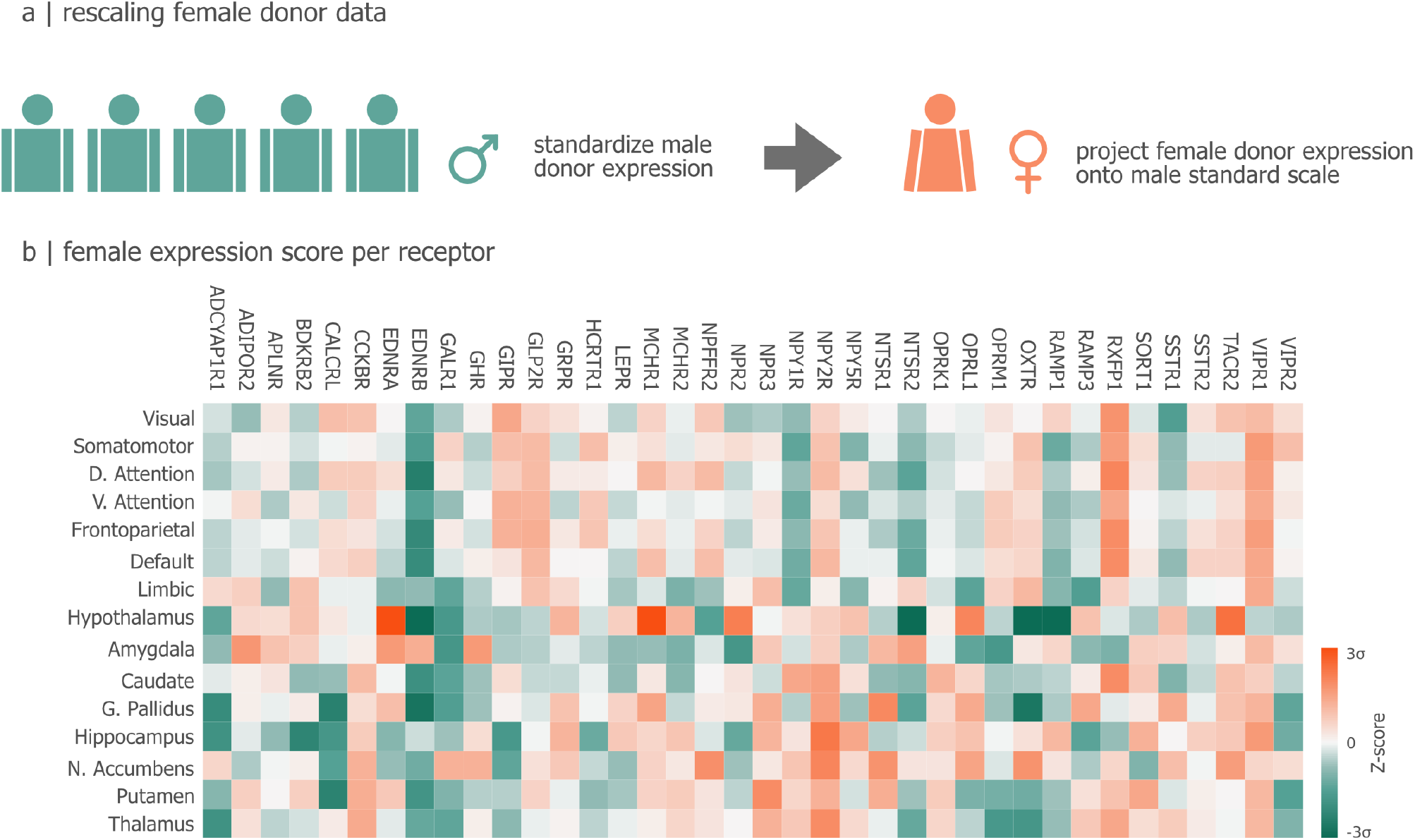
Sex differences in neuropeptide receptor expression. (a) To identify receptors with differential expression between the single female donor and the remaining five male donors, we z-score the female donor data with respect to male donor expression. (b) The resulting gene distribution highlights neuropeptide receptor genes with greater (orange) or lower (green) differential expression in the female donor.

**Figure S3.**
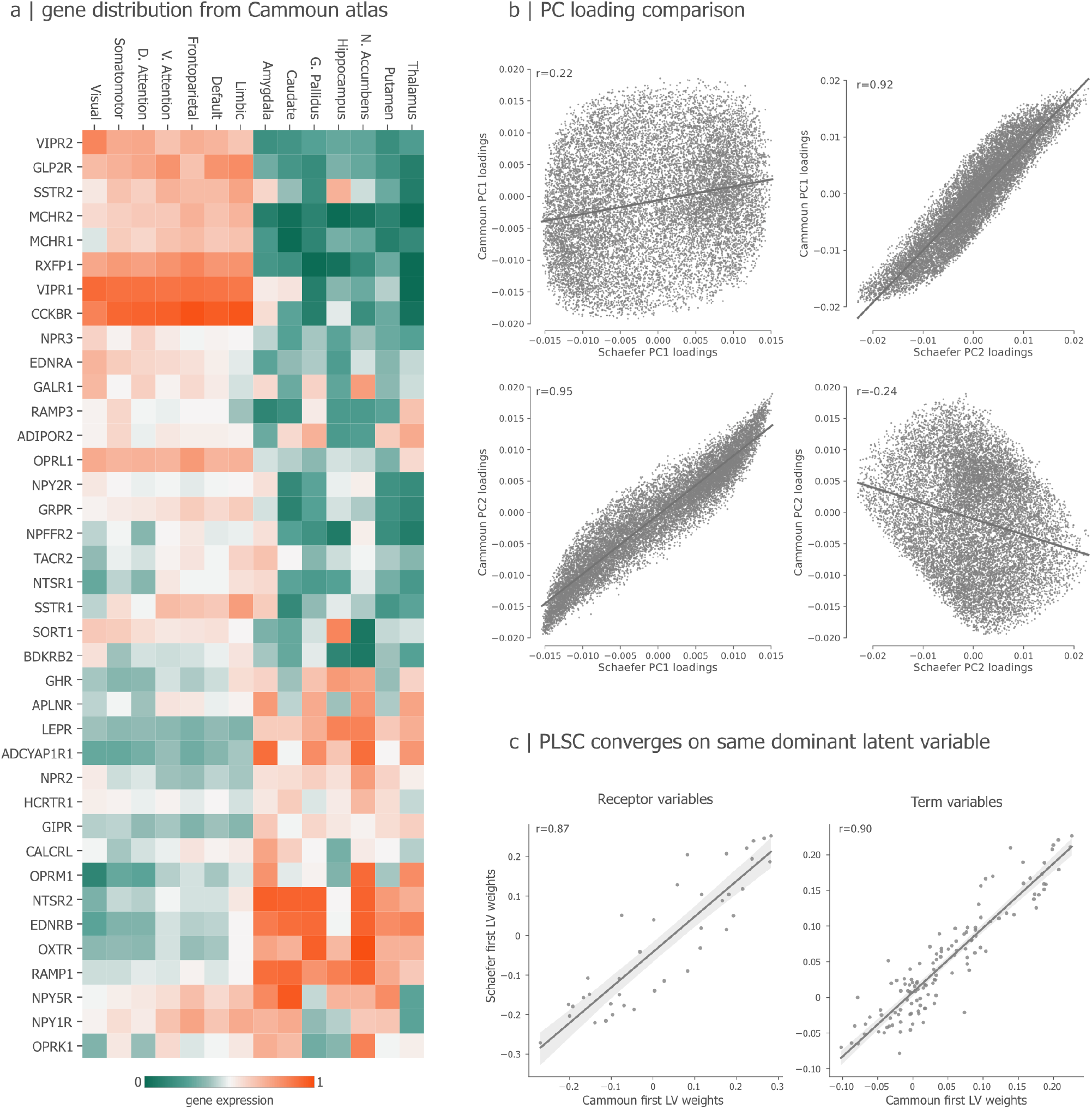
Replication with an anatomical landmark-based atlas. Main text figures are all shown for a functionally-derived atlas of the cortex and subcortex [147, 167]. Here we re-parcellate the cortical data according the Cammoun atlas [25], a subdivision of the landmark-based Desikan-Killiany atlas [35], yielding 500 regions of interest. We apply a corresponding FreeSurfer anatomical parcellation to the subcortex [48]. (a) Gene expression matrix of neuropeptide receptors, organized identically to Fig. 2. (b) PC1 and PC2 of the functional and anatomical parcellations. Collectively, the two PCs account for approximately equal portions of variance (PC1=16.58%, PC2=15.38%), but have reversed order. Namely, anatomical PC1 closely matches functional PC2 (*r* = 0.92, *p* = 0), and anatomical PC2 closely matches functional PC1 (*r* = 0.95, *p* = 0). (c) To ensure this reversal does not affect our covariance-based PLSC analysis in Fig. 5, we repeat the analysis using the anatomical parcellation. The scatterplots show positive correspondence between the PLSC weights (singular vectors) derived using the anatomical and functional parcellations(*r* = 0.87,*p* = 10^−12^);term weight similarity *r* = 0.9,*p* = 6 *×* 10^−47^.

**Figure S4.**
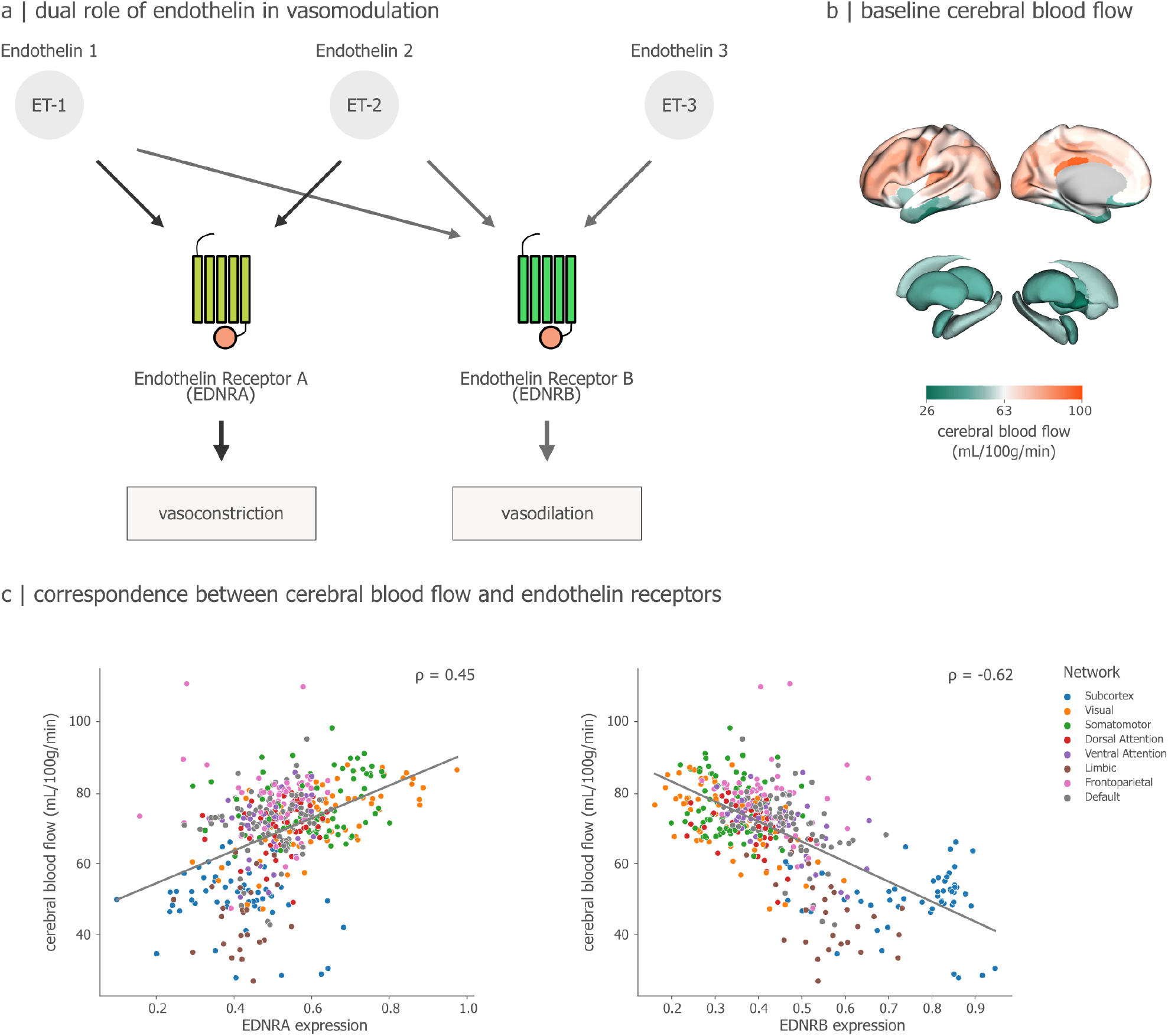
Vasomodulatory neuropeptides correlate with cerebral blood flow. To confirm specificity of neuropeptide mapping, we assess whether regional differences in the vasomodulatory endothelin receptors correspond to empirical measurements of cerebral blood flow, as measured using arterial spin labeling (ASL; see *Methods*). (a) We focus on two endothelin receptors, vasoconstrictive endothelin receptor A (EDNRA) and vasodilutive endothelin receptor B (EDNRB). (b) We use a group average template of cerebral blood flow from arterial spin labeling acquisitions of n=678 individuals [69]. (c) Regional expression of vasoconstrictive EDNRA is positively correlated with cerebral blood flow (*ρ* = 0.45, *p*_SMASH_ = 0.0011), while regional expression of vasodilutive EDNRB is negatively correlated with cerebral blood flow (*ρ* = −0.62, *p*_SMASH_ = 0.0001). This is consistent with the regulatory role of endothelin peptides, responding to blood flow changes through extensive constriction in high perfusion and dilation in low perfusion areas, thereby keeping blood supply within physiological ranges [4, 91, 105, 117].

**Figure S5.**
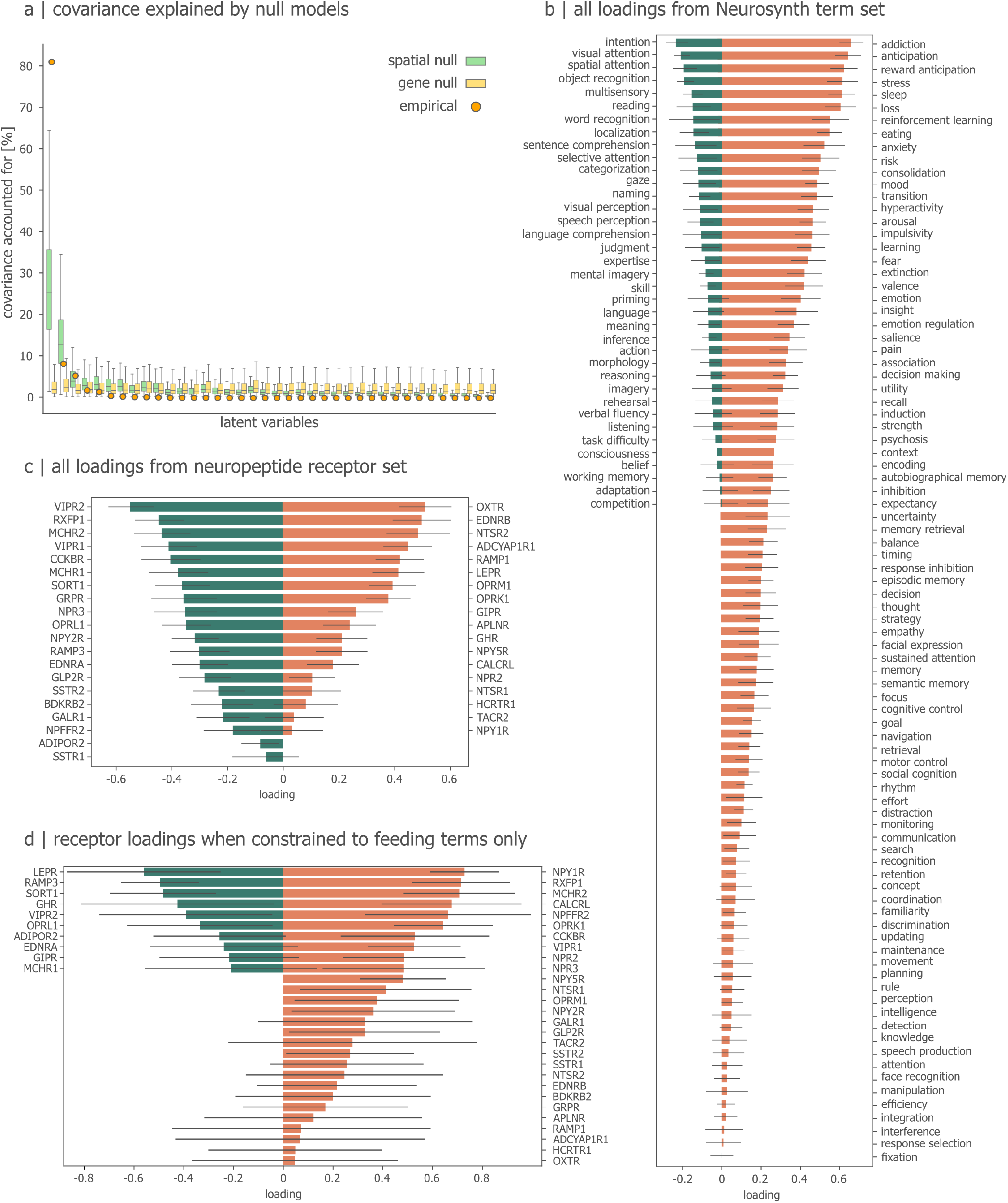
Supporting information for the PLSC model. (a) The covariance explained by each latent variable (orange dots) shown alongside null distributions generated using spatial autocorrelation-preserving randomization (green boxplots) and matched null genes (yellow boxplots) (see *Methods*). (b) The full set of loadings for all Neurosynth terms in the Cognitive Atlas. (c) The full set of loadings for all neuropeptide receptors. (d) Loadings for neuropeptide receptors when the analysis is constrained only to feeding terms.

**Figure S6.**
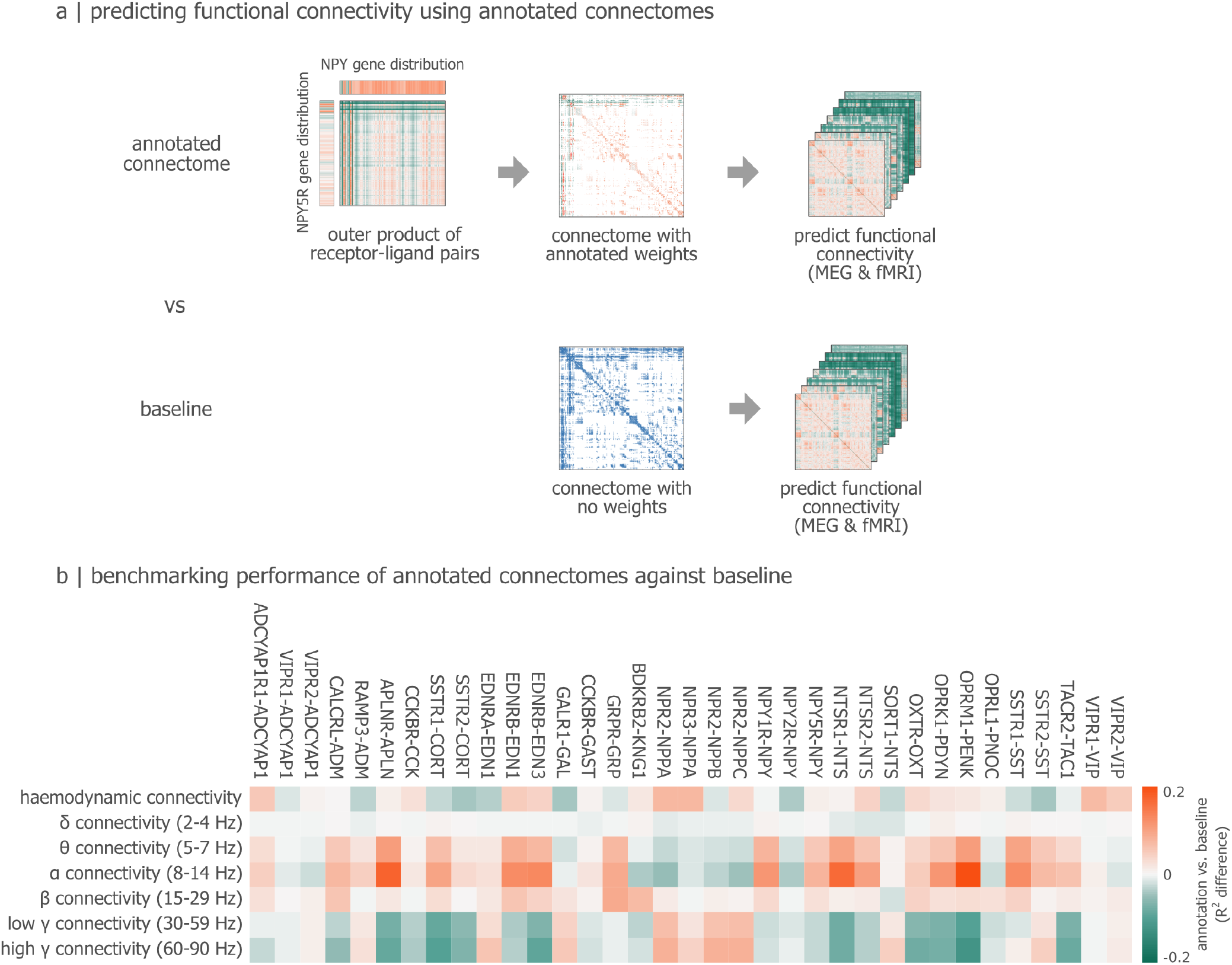
Benchmarking peptide annotated connectomes on all connectivity modes. (a) We re-weigh dMRI-derived structural connectivity networks using neuropeptide receptor-ligand correspondence (see *Methods*). We then use these neuropeptide-weighted connectomes to predict MEG- and fMRI-derived functional connectivity networks. Results are expressed relative to baseline, in which the original dMRI matrix is used without neuroepetide weights. (b) Changes in structure-function coupling when applying neuropeptide weights, shown for multiple receptor-ligand pairs (columns) and multiple modes of functional connectivity (rows).

## References

[1] Abraham, W. C. and Bear, M. F. (1996). Metaplasticity: the plasticity of synaptic plasticity. Trends in neurosciences, 19(4):126–130.

[2] Alonso, A., Faure, M.-P., and Beaudet, A. (1994). Neurotensin promotes oscillatory bursting behavior and is internalized in basal forebrain cholinergic neurons. Journal of Neuroscience, 14(10):5778–5792.

[3] Arnatkeviciūtė, A., Fulcher, B. D., and Fornito, A. (2019). A practical guide to linking brain-wide gene expression and neuroimaging data. Neuroimage, 189:353–367.

[4] Ashby, J. W. and Mack, J. J. (2021). Endothelial control of cerebral blood flow. The American Journal of Pathology, 191(11):1906–1916.

[5] Avena-Koenigsberger, A., Misic, B., and Sporns, O. (2018). Communication dynamics in complex brain networks. Nature reviews neuroscience, 19(1):17–33.

[6] Azen, R. and Budescu, D. V. (2003). The dominance analysis approach for comparing predictors in multiple regression. Psychological methods, 8(2):129.

[7] Bale, T. L. and Dorsa, D. M. (1995). Sex differences in and effects of estrogen on oxytocin receptor messenger ribonucleic acid expression in the ventromedial hypothalamus. Endocrinology, 136(1):27–32.

[8] Barbas, H. (2015). General cortical and special prefrontal connections: principles from structure to function. Annual review of neuroscience, 38:269–289.

[9] Bartz, J. A., Zaki, J., Bolger, N., and Ochsner, K. N. (2011). Social effects of oxytocin in humans: context and person matter. Trends in cognitive sciences, 15(7):301–309.

[10] Bates, S. H. and Myers, M. G. (2003). The role of leptin receptor signaling in feeding and neuroendocrine function. Trends in Endocrinology & Metabolism, 14(10):447– 452.

[11] Bazinet, V., de Wael, R. V., Hagmann, P., Bernhardt, B. C., and Misic, B. (2021). Multiscale communication in cortico-cortical networks. NeuroImage, 243:118546.

[12] Bazinet, V., Hansen, J. Y., and Misic, B. (2023a). Towards a biologically annotated brain connectome. Nature reviews neuroscience, 24(12):747–760.

[13] Bazinet, V., Hansen, J. Y., Vos de Wael, R., Bernhardt, B. C., van den Heuvel, M. P., and Misic, B. (2023b). Assortative mixing in micro-architecturally annotated brain connectomes. Nature Communications, 14(1):2850.

[14] Beliveau, V., Ganz, M., Feng, L., Ozenne, B., Højgaard, L., Fisher, P. M., Svarer, C., Greve, D. N., and Knudsen, G. M. (2017). A high-resolution in vivo atlas of the human brain’s serotonin system. Journal of Neuroscience, 37(1):120–128.

[15] Betzel, R. F., Griffa, A., Avena-Koenigsberger, A., Goñi, J., Thiran, J.-P., Hagmann, P., and Sporns, O. (2013). Multi-scale community organization of the human structural connectome and its relationship with resting-state functional connectivity. Network Science, 1(3):353–373.

[16] Betzel, R. F., Griffa, A., Hagmann, P., and Mišić, B. (2019). Distance-dependent consensus thresholds for generating group-representative structural brain networks. Network neuroscience, 3(2):475–496.

[17] Beul, S. F., Goulas, A., and Hilgetag, C. C. (2021). An architectonic type principle in the development of laminar patterns of cortico-cortical connections. Brain Structure and Function, 226(4):979–987.

[18] Beul, S. F. and Hilgetag, C. C. (2020). Systematic modelling of the development of laminar projection origins in the cerebral cortex: Interactions of spatiotemporal patterns of neurogenesis and cellular heterogeneity. PLoS Computational Biology, 16(10):e1007991.

[19] Bhat, U. S., Shahi, N., Surendran, S., and Babu, K. (2021). Neuropeptides and behaviors: how small peptides regulate nervous system function and behavioral outputs. Frontiers in Molecular Neuroscience, 14:786471.

[20] Bruns, A., Eckhorn, R., Jokeit, H., and Ebner, A. (2000). Amplitude envelope correlation detects coupling among incoherent brain signals. Neuroreport, 11(7):1509– 1514.

[21] Burbach, J. P. H. (2010). Neuropeptides from concept to online database www.neuropeptides.nl. European journal of pharmacology, 626(1):27–48.

[22] Burbach, J. P. H. (2011). What are neuropeptides? Neuropeptides: Methods and protocols, pages 1–36.

[23] Burt, J. B., Demirtas, M., Eckner, W. J., Navejar, N. M., Ji, J. L., Martin, W. J., Bernacchia, A., Anticevic, A., and Murray, J. D. (2018). Hierarchy of transcriptomic specialization across human cortex captured by structural neuroimaging topography. Nature neuroscience, 21(9):1251–1259.

[24] Burt, J. B., Helmer, M., Shinn, M., Anticevic, A., and Murray, J. D. (2020). Generative modeling of brain maps with spatial autocorrelation. NeuroImage, 220:117038.

[25] Cammoun, L., Gigandet, X., Meskaldji, D., Thiran, J. P., Sporns, O., Do, K. Q., Maeder, P., Meuli, R., and Hagmann, P. (2012). Mapping the human connectome at multiple scales with diffusion spectrum mri. Journal of neuroscience methods, 203(2):386–397.

[26] Cape, E. G., Manns, I. D., Alonso, A., Beaudet, A., and Jones, B. E. (2000). Neurotensin-induced bursting of cholinergic basal forebrain neurons promotes γ and θ cortical activity together with waking and paradoxical sleep. Journal of Neuroscience, 20(22):8452–8461.

[27] Cauzzo, S., Singh, K., Stauder, M., García-Gomar, M. G., Vanello, N., Passino, C., Staab, J., Indovina, I., and Bianciardi, M. (2022). Functional connectome of brainstem nuclei involved in autonomic, limbic, pain and sensory processing in living humans from 7 tesla resting state fmri. NeuroImage, 250:118925.

[28] Chen, M., Yan, H.-h., Shu, S., Pei, L., Zang, L.-k., Fu, Y., Wang, Z.-f., Wan, Q., and Bi, L.-l. (2017). Amygdalar endothelin-1 regulates pyramidal neuron excitability and affects anxiety. Scientific Reports, 7(1):2316.

[29] Cooper, S., Robison, A., and Mazei-Robison, M. S. (2017). Reward circuitry in addiction. Neurotherapeutics, 14(3):687–697.

[30] Crown, A., Clifton, D. K., and Steiner, R. A. (2007). Neuropeptide signaling in the integration of metabolism and reproduction. Neuroendocrinology, 86(3):175–182.

[31] Darcq, E. and Kieffer, B. L. (2018). Opioid receptors: drivers to addiction? Nature Reviews Neuroscience, 19(8):499–514.

[32] Dashwood, M. R. and Loesch, A. (2010). Endothelin-1 as a neuropeptide: neurotransmitter or neurovascular effects? Journal of cell communication and signaling, 4:51–62.

[33] Dear, R., Wagstyl, K., Seidlitz, J., Markello, R. D., Arnatkevičiūtė, A., Anderson, K. M., Bethlehem, R. A., Consortium, L. B. C., Raznahan, A., Bullmore, E. T., et al. (2024). Cortical gene expression architecture links healthy neurodevelopment to the imaging, transcriptomics and genetics of autism and schizophrenia. Nature Neuroscience, pages 1–12.

[34] Dehghan, F., Muniandy, S., Yusof, A., and Salleh, N. (2014). Sex-steroid regulation of relaxin receptor isoforms (rxfp1 & rxfp2) expression in the patellar tendon and lateral collateral ligament of female wky rats. International journal of medical sciences, 11(2):180.

[35] Desikan, R. S., Ségonne, F., Fischl, B., Quinn, B. T., Dickerson, B. C., Blacker, D., Buckner, R. L., Dale, A. M., Maguire, R. P., Hyman, B. T., et al. (2006). An automated labeling system for subdividing the human cerebral cortex on mri scans into gyral based regions of interest. Neuroimage, 31(3):968–980.

[36] Destrieux, C., Fischl, B., Dale, A., and Halgren, E. (2010). Automatic parcellation of human cortical gyri and sulci using standard anatomical nomenclature. Neuroimage, 53(1):1–15.

[37] Detre, J. A., Zhang, W., Roberts, D. A., Silva, A. C., Williams, D. S., Grandis, D. J., Koretsky, A. P., and Leigh, J. S. (1994). Tissue specific perfusion imaging using arterial spin labeling. NMR in Biomedicine, 7(1-2):75–82.

[38] DuBois, J. M., Rousset, O. G., Rowley, J., Porras-Betancourt, M., Reader, A. J., Labbe, A., Massarweh, G., Soucy, J.-P., Rosa-Neto, P., and Kobayashi, E. (2016). Characterization of age/sex and the regional distribution of mglur5 availability in the healthy human brain measured by high-resolution [11 c] abp688 pet. European journal of nuclear medicine and molecular imaging, 43:152–162.

[39] Edgar, R. C. (2004). Muscle: multiple sequence alignment with high accuracy and high throughput. Nucleic acids research, 32(5):1792–1797.

[40] Egerton, A., Mehta, M. A., Montgomery, A. J., Lappin, J. M., Howes, O. D., Reeves, S. J., Cunningham, V. J., and Grasby, P. M. (2009). The dopaminergic basis of human behaviors: A review of molecular imaging studies. Neuroscience & Biobehavioral Reviews, 33(7):1109–1132.

[41] Eickhoff, S. B., Bzdok, D., Laird, A. R., Kurth, F., and Fox, P. T. (2012). Activation likelihood estimation meta-analysis revisited. Neuroimage, 59(3):2349–2361.

[42] Eiden, L. E., Hernández, V. S., Jiang, S. Z., and Zhang, L. (2022). Neuropeptides and small-molecule amine transmitters: cooperative signaling in the nervous system. Cellular and Molecular Life Sciences, 79(9):492.

[43] Elam, J. S., Glasser, M. F., Harms, M. P., Sotiropoulos, S. N., Andersson, J. L., Burgess, G. C., Curtiss, S. W., Oostenveld, R., Larson-Prior, L. J., Schoffelen, J.-M., et al. (2021). The human connectome project: a retrospective. NeuroImage, 244:118543.

[44] Elmquist, J. K., Elias, C. F., and Saper, C. B. (1999). From lesions to leptin: hypothalamic control of food intake and body weight. Neuron, 22(2):221–232.

[45] Farde, L., Pauli, S., Hall, H., Eriksson, L., Halldin, C., Högberg, T., Nilsson, L., Sjögren, I., and Stone-Elander, S. (1988). Stereoselective binding of 11c-raclopride in living human brain—a search for extrastriatal central d2-dopamine receptors by pet. Psychopharmacology, 94:471–478.

[46] Federhen, S. (2012). The ncbi taxonomy database. Nucleic acids research, 40(D1):D136–D143.

[47] Figley, C. R., Uddin, M. N., Wong, K., Kornelsen, J., Puig, J., and Figley, T. D. (2022). Potential pitfalls of using fractional anisotropy, axial diffusivity, and radial diffusivity as biomarkers of cerebral white matter microstructure. Frontiers in Neuroscience, 15:799576.

[48] Fischl, B., Salat, D. H., Busa, E., Albert, M., Dieterich, M., Haselgrove, C., Van Der Kouwe, A., Killiany, R., Kennedy, D., Klaveness, S., et al. (2002). Whole brain segmentation: automated labeling of neuroanatomical structures in the human brain. Neuron, 33(3):341–355.

[49] Fox, A. S., Chang, L. J., Gorgolewski, K. J., and Yarkoni, T. (2014). Bridging psychology and genetics using largescale spatial analysis of neuroimaging and neurogenetic data. bioRxiv, page 012310.

[50] Frémaux, N. and Gerstner, W. (2016). Neuromodulated spike-timing-dependent plasticity, and theory of three-factor learning rules. Frontiers in neural circuits, 9:85.

[51] Friedman, M. (1940). A comparison of alternative tests of significance for the problem of m rankings. The annals of mathematical statistics, 11(1):86–92.

[52] Froudist-Walsh, S., Xu, T., Niu, M., Rapan, L., Zhao, L., Margulies, D. S., Zilles, K., Wang, X.-J., and Palomero-Gallagher, N. (2023). Gradients of neurotransmitter receptor expression in the macaque cortex. Nature neuroscience, 26(7):1281–1294.

[53] Fulcher, B. D. and Fornito, A. (2016). A transcriptional signature of hub connectivity in the mouse connectome. Proceedings of the National Academy of Sciences, 113(5):1435–1440.

[54] Gallezot, J.-D., Nabulsi, N., Neumeister, A., Planeta-Wilson, B., Williams, W. A., Singhal, T., Kim, S., Maguire, R. P., McCarthy, T., Frost, J. J., et al. (2010). Kinetic modeling of the serotonin 5-ht1b receptor radioligand [11c] p943 in humans. Journal of Cerebral Blood Flow & Metabolism, 30(1):196–210.

[55] Gallezot, J.-D., Planeta, B., Nabulsi, N., Palumbo, D., Li, X., Liu, J., Rowinski, C., Chidsey, K., Labaree, D., Ropchan, J., et al. (2017). Determination of receptor occupancy in the presence of mass dose:[11c] gsk189254 pet imaging of histamine h3 receptor occupancy by pf03654746. Journal of Cerebral Blood Flow & Metabolism, 37(3):1095–1107.

[56] García-Cabezas, M.Á., Zikopoulos, B., and Barbas, H. (2019). The structural model: a theory linking connections, plasticity, pathology, development and evolution of the cerebral cortex. Brain Structure and Function, 224(3):985–1008.

[57] Gardner, E. L. (2011). Addiction and brain reward and antireward pathways. Chronic Pain and Addiction, 30:22–60.

[58] Gilden, L. and Kozakiewicz, R. (1976). The effect of morphine on the eeg of the hypothalamus in the rat. Physiology & Behavior, 16(2):169–176.

[59] Glasser, M. F., Sotiropoulos, S. N., Wilson, J. A., Coalson, T. S., Fischl, B., Andersson, J. L., Xu, J., Jbabdi, S., Webster, M., Polimeni, J. R., et al. (2013). The minimal preprocessing pipelines for the human connectome project. Neuroimage, 80:105–124.

[60] Goñi, J., Van Den Heuvel, M. P., Avena-Koenigsberger, A., Velez de Mendizabal, N., Betzel, R. F., Griffa, A., Hagmann, P., Corominas-Murtra, B., Thiran, J.-P., and Sporns, O. (2014). Resting-brain functional connectivity predicted by analytic measures of network communication. Proc Natl Acad Sci USA, 111(2):833–838.

[61] Gorka, S. M., Fitzgerald, D. A., De Wit, H., Angstadt, M., and Phan, K. L. (2014). Opioid modulation of restingstate anterior cingulate cortex functional connectivity. Journal of Psychopharmacology, 28(12):1115–1124.

[62] Goulas, A., Margulies, D. S., Bezgin, G., and Hilgetag, C. C. (2019). The architecture of mammalian cortical connectomes in light of the theory of the dual origin of the cerebral cortex. Cortex, 118:244–261.

[63] Goulas, A., Zilles, K., and Hilgetag, C. C. (2018). Cortical gradients and laminar projections in mammals. Trends in neurosciences, 41(11):775–788.

[64] Hansen, J. Y., Cauzzo, S., Singh, K., García-Gomar, M. G., Shine, J. M., Bianciardi, M., and Misic, B. (2024). Integrating brainstem and cortical functional architectures. Nature Neuroscience, pages 1–12.

[65] Hansen, J. Y., Markello, R. D., Tuominen, L., Nørgaard, M., Kuzmin, E., Palomero-Gallagher, N., Dagher, A., and Misic, B. (2022a). Correspondence between gene expression and neurotransmitter receptor and transporter density in the human brain. NeuroImage, 264:119671.

[66] Hansen, J. Y., Markello, R. D., Vogel, J. W., Seidlitz, J., Bzdok, D., and Misic, B. (2021). Mapping gene transcription and neurocognition across human neocortex. Nature Human Behaviour, 5(9):1240–1250.

[67] Hansen, J. Y., Shafiei, G., Markello, R. D., Smart, K., Cox, S. M., Nørgaard, M., Beliveau, V., Wu, Y., Gallezot, J.-D., Aumont, É., et al. (2022b). Mapping neurotransmitter systems to the structural and functional organization of the human neocortex. Nature Neuroscience, pages 1–13.

[68] Hansen, J. Y., Shafiei, G., Voigt, K., Liang, E. X., Cox, S. M., Leyton, M., Jamadar, S. D., and Misic, B. (2023). Integrating multimodal and multiscale connectivity blueprints of the human cerebral cortex in health and disease. Plos Biology, 21(9):e3002314.

[69] Harms, M. P., Somerville, L. H., Ances, B. M., Andersson, J., Barch, D. M., Bastiani, M., Bookheimer, S. Y., Brown, T. B., Buckner, R. L., Burgess, G. C., et al. (2018). Extending the human connectome project across ages: Imaging protocols for the lifespan development and aging projects. Neuroimage, 183:972–984.

[70] Hawrylycz, M., Miller, J. A., Menon, V., Feng, D., Dolbeare, T., Guillozet-Bongaarts, A. L., Jegga, A. G., Aronow, B. J., Lee, C.-K., Bernard, A., et al. (2015). Canonical genetic signatures of the adult human brain. Nature neuroscience, 18(12):1832–1844.

[71] Hawrylycz, M. J., Lein, E. S., Guillozet-Bongaarts, A. L., Shen, E. H., Ng, L., Miller, J. A., Van De Lagemaat, L. N., Smith, K. A., Ebbert, A., Riley, Z. L., et al. (2012). An anatomically comprehensive atlas of the adult human brain transcriptome. Nature, 489(7416):391–399.

[72] Heasley, L. E. (2001). Autocrine and paracrine signaling through neuropeptide receptors in human cancer. Oncogene, 20(13):1563–1569.

[73] Hillmer, A. T., Esterlis, I., Gallezot, J.-D., Bois, F., Zheng, M.-Q., Nabulsi, N., Lin, S.-F., Papke, R., Huang, Y., Sabri, O., et al. (2016). Imaging of cerebral α4β2* nicotinic acetylcholine receptors with (-)-[18f] flubatine pet: Implementation of bolus plus constant infusion and sensitivity to acetylcholine in human brain. Neuroimage, 141:71–80.

[74] Hökfelt, T., Broberger, C., Xu, Z.-Q. D., Sergeyev, V., Ubink, R., and Diez, M. (2000a). Neuropeptides—an overview. Neuropharmacology, 39(8):1337–1356.

[75] Hökfelt, T., Broberger, C., Xu, Z.-Q. D., Sergeyev, V., Ubink, R., and Diez, M. (2000b). Neuropeptides—an overview. Neuropharmacology, 39(8):1337–1356.

[76] Huang, P., Xiang, X., Chen, X., and Li, H. (2020). Somatostatin neurons govern theta oscillations induced by salient visual signals. Cell Reports, 33(8).

[77] Huntenburg, J. M., Bazin, P.-L., Goulas, A., Tardif, C. L., Villringer, A., and Margulies, D. S. (2017). A systematic relationship between functional connectivity and intracortical myelin in the human cerebral cortex. Cerebral Cortex, 27(2):981–997.

[78] Ivell, R., Agoulnik, A. I., and Anand-Ivell, R. (2017). Relaxin-like peptides in male reproduction–a human perspective. British Journal of Pharmacology, 174(10):990–1001.

[79] Ji, J. L., Demšar, J., Fonteneau, C., Tamayo, Z., Pan, L., Kraljič, A., Matkovič, A., Purg, N., Helmer, M., Warrington, S., et al. (2023). Qunex—an integrative platform for reproducible neuroimaging analytics. Frontiers in Neuroinformatics, 17:1104508.

[80] Karalija, N., Jonassson, L., Johansson, J., Papenberg, G., Salami, A., Andersson, M., Riklund, K., Nyberg, L., and Boraxbekk, C.-J. (2020). High long-term test– retest reliability for extrastriatal 11c-raclopride binding in healthy older adults. Journal of Cerebral Blood Flow & Metabolism, 40(9):1859–1868.

[81] Kendall, M. G. and Smith, B. B. (1939). The problem of m rankings. The annals of mathematical statistics, 10(3):275–287.

[82] Kirk, T. F., McConnell, F. A. K., Toner, J., Craig, M. S., Carone, D., Li, X., Suzuki, Y., Coalson, T. S., Harms, M. P., Glasser, M. F., et al. (2024). Arterial spin labelling perfusion mri analysis for the human connectome project lifespan ageing and development studies. bioRxiv, pages 2024–09.

[83] Klok, M. D., Jakobsdottir, S., and Drent, M. L. (2007). The role of leptin and ghrelin in the regulation of food intake and body weight in humans: a review. Obesity reviews, 8(1):21–34.

[84] Knutson, B., Adams, C. M., Fong, G. W., and Hommer, D. (2001). Anticipation of increasing monetary reward selectively recruits nucleus accumbens. The Journal of neuroscience, 21(16):RC159.

[85] Kosakovsky Pond, S. L., Murrell, B., Fourment, M., Frost, S. D., Delport, W., and Scheffler, K. (2011). A random effects branch-site model for detecting episodic diversifying selection. Molecular biology and evolution, 28(11):3033–3043.

[86] Kosakovsky Pond, S. L., Poon, A. F., Velazquez, R., Weaver, S., Hepler, N. L., Murrell, B., Shank, S. D., Magalis, B. R., Bouvier, D., Nekrutenko, A., et al. (2020). Hyphy 2.5—a customizable platform for evolutionary hypothesis testing using phylogenies. Molecular biology and evolution, 37(1):295–299.

[87] Krahn, D., Gosnell, B. A., Levine, A. S., and Morley, J. E. (1984). Effects of calcitonin gene-related peptide on food intake. Peptides, 5(5):861–864.

[88] Kumar, S., Stecher, G., Suleski, M., and Hedges, S. B. (2017). Timetree: a resource for timelines, timetrees, and divergence times. Molecular biology and evolution, 34(7):1812–1819.

[89] Larsson, H., Elmståhl, S., Berglund, G., and Ahrén, B. (1998). Evidence for leptin regulation of food intake in humans. The Journal of Clinical Endocrinology & Metabolism, 83(12):4382–4385.

[90] Leuker, C., Pachur, T., Hertwig, R., and Pleskac, T. J. (2018). Exploiting risk–reward structures in decision making under uncertainty. Cognition, 175:186–200.

[91] Li, T., Li, D., Wei, Q., Shi, M., Xiang, J., Gao, R., Chen, C., and Xu, Z.-X. (2023). Dissecting the neurovascular unit in physiology and alzheimer’s disease: functions, imaging tools and genetic mouse models. Neurobiology of Disease, 181:106114.

[92] Lin, L., Faraco, J., Li, R., Kadotani, H., Rogers, W., Lin, X., Qiu, X., de Jong, P. J., Nishino, S., and Mignot, E. (1999). The sleep disorder canine narcolepsy is caused by a mutation in the hypocretin (orexin) receptor 2 gene. Cell, 98(3):365–376.

[93] Liu, D., Rahman, M., Johnson, A., Tsutsui-Kimura, I., Pena, N., Talay, M., Logeman, B. L., Finkbeiner, S., Choi, S., Capo-Battaglia, A., et al. (2023a). A hypothalamic circuit underlying the dynamic control of social homeostasis. BioRxiv.

[94] Liu, Z.-Q., Shafiei, G., Baillet, S., and Misic, B. (2023b). Spatially heterogeneous structure-function coupling in haemodynamic and electromagnetic brain networks. NeuroImage, 278:120276.

[95] Luppi, A. I., Liu, Z.-Q., Hansen, J. Y., Cofre, R., Kuzmin, E., Froudist-Walsh, S., Palomero-Gallagher, N., and Misic, B. (2024). Benchmarking macaque brain gene expression for horizontal and vertical translation. bioRxiv, pages 2024–08.

[96] Maier-Hein, K. H., Neher, P. F., Houde, J.-C., Côté, M.-A., Garyfallidis, E., Zhong, J., Chamberland, M., Yeh, F.-C., Lin, Y.-C., Ji, Q., et al. (2017). The challenge of mapping the human connectome based on diffusion tractography. Nature communications, 8(1):1349.

[97] Marder, E. (2012). Neuromodulation of neuronal circuits: back to the future. Neuron, 76(1):1–11.

[98] Markello, R. D., Arnatkeviciute, A., Poline, J.-B., Fulcher, B. D., Fornito, A., and Misic, B. (2021). Standardizing workflows in imaging transcriptomics with the abagen toolbox. elife, 10:e72129.

[99] Markello, R. D., Hansen, J. Y., Liu, Z.-Q., Bazinet, V., Shafiei, G., Suárez, L. E., Blostein, N., Seidlitz, J., Baillet, S., Satterthwaite, T. D., et al. (2022). Neuromaps: structural and functional interpretation of brain maps. Nature Methods, 19(11):1472–1479.

[100] Markello, R. D. and Misic, B. (2021). Comparing spatial null models for brain maps. NeuroImage, page 118052.

[101] Martin, J., Renaud, L., and Brazeau, P. (1975). Hypothalamic peptides: New evidence for” peptidergic” pathways in the cns. The Lancet, 306(7931):393–395.

[102] McIntosh, A. R. and Lobaugh, N. J. (2004). Partial least squares analysis of neuroimaging data: applications and advances. Neuroimage, 23:S250–S263.

[103] McIntosh, A. R. and Mišić, B. (2013). Multivariate statistical analyses for neuroimaging data. Annu Rev Psychol, 64(1):499–525.

[104] Meister, B. (2000). Control of food intake via leptin receptors in the hypothalamus.

[105] Michinaga, S., Hishinuma, S., and Koyama, Y. (2023). Roles of astrocytic endothelin etb receptor in traumatic brain injury. Cells, 12(5):719.

[106] Miller, J. A., Menon, V., Goldy, J., Kaykas, A., Lee, C.-K., Smith, K. A., Shen, E. H., Phillips, J. W., Lein, E. S., and Hawrylycz, M. J. (2014). Improving reliability and absolute quantification of human brain microarray data by filtering and scaling probes using rna-seq. BMC genomics, 15:1–14.

[107] Mirchi, N., Betzel, R. F., Bernhardt, B. C., Dagher, A., and Mišić, B. (2019). Tracking mood fluctuations with functional network patterns. Soc Cogn Affect Neurosci, 14(1):47–57.

[108] Naganawa, M., Nabulsi, N., Henry, S., Matuskey, D., Lin, S.-F., Slieker, L., Schwarz, A. J., Kant, N., Jesudason, C., Ruley, K., et al. (2021). First-in-human assessment of 11c-lsn3172176, an m1 muscarinic acetylcholine receptor pet radiotracer. Journal of Nuclear Medicine, 62(4):553–560.

[109] Nakagawa, Y., Nishikimi, T., and Kuwahara, K. (2019). Atrial and brain natriuretic peptides: Hormones secreted from the heart. Peptides, 111:18–25.

[110] Nasseef, M. T., Singh, J. P., Ehrlich, A. T., McNicholas, M., Park, D. W., Ma, W., Kulkarni, P., Kieffer, B. L., and Darcq, E. (2019). Oxycodone-mediated activation of the mu opioid receptor reduces whole brain functional connectivity in mice. ACS Pharmacology & Translational Science, 2(4):264–274.

[111] Naufahu, J., Cunliffe, A. D., and Murray, J. F. (2013). The roles of melanin-concentrating hormone in energy balance and reproductive function: are they connected? Reproduction, 146(5):R141–R150.

[112] Nestler, E. J. and Carlezon Jr, W. A. (2006). The mesolimbic dopamine reward circuit in depression. Biological psychiatry, 59(12):1151–1159.

[113] Nørgaard, M., Beliveau, V., Ganz, M., Svarer, C., Pinborg, L. H., Keller, S. H., Jensen, P. S., Greve, D. N., and Knudsen, G. M. (2021). A high-resolution in vivo atlas of the human brain’s benzodiazepine binding site of gabaa receptors. NeuroImage, 232:117878.

[114] Normandin, M. D., Zheng, M.-Q., Lin, K.-S., Mason, N. S., Lin, S.-F., Ropchan, J., Labaree, D., Henry, S., Williams, W. A., Carson, R. E., et al. (2015). Imaging the cannabinoid cb1 receptor in humans with [11c] omar: assessment of kinetic analysis methods, test–retest reproducibility, and gender differences. Journal of Cerebral Blood Flow & Metabolism, 35(8):1313–1322.

[115] Nusbaum, M. P. and Blitz, D. M. (2012). Neuropeptide modulation of microcircuits. Current opinion in neurobiology, 22(4):592–601.

[116] Nusbaum, M. P., Blitz, D. M., and Marder, E. (2017). Functional consequences of neuropeptide and smallmolecule co-transmission. Nature Reviews Neuroscience, 18(7):389–403.

[117] Pacholko, A. and Iadecola, C. (2024). Hypertension, neurodegeneration, and cognitive decline. Hypertension, 81(5):991–1007.

[118] Pauli, W. M., Nili, A. N., and Tyszka, J. M. (2018). A high-resolution probabilistic in vivo atlas of human subcortical brain nuclei. Scientific data, 5(1):1–13.

[119] Pearson, W. R. (2013). An introduction to sequence similarity (“homology”) searching. Current protocols in bioinformatics, 42(1):3–1.

[120] Peineau, S., Rabiant, K., Pierrefiche, O., and Potier, B. (2018). Synaptic plasticity modulation by circulating peptides and metaplasticity: involvement in alzheimer’s disease. Pharmacological Research, 130:385–401.

[121] Pénicaud, L., Meillon, S., and Brondel, L. (2012). Leptin and the central control of feeding behavior. Biochimie, 94(10):2069–2074.

[122] Perlow, M. J., Freed, W. J., Carman, J. S., and Wyatt, R. J. (1980). Calcitonin reduces feeding in man, monkey and rat. Pharmacology Biochemistry and Behavior, 12(4):609–612.

[123] Pfeffer, C. K., Xue, M., He, M., Huang, Z. J., and Scanziani, M. (2013). Inhibition of inhibition in visual cortex: the logic of connections between molecularly distinct interneurons. Nature neuroscience, 16(8):1068– 1076.

[124] Pi, H.-J., Hangya, B., Kvitsiani, D., Sanders, J. I., Huang, Z. J., and Kepecs, A. (2013). Cortical interneurons that specialize in disinhibitory control. Nature, 503(7477):521–524.

[125] Pizza, F., Barateau, L., Dauvilliers, Y., and Plazzi, G. (2022). The orexin story, sleep and sleep disturbances. Journal of Sleep Research, 31(4):e13665.

[126] Poldrack, R. A., Kittur, A., Kalar, D., Miller, E., Seppa, C., Gil, Y., Parker, D. S., Sabb, F. W., and Bilder, R. M. (2011). The cognitive atlas: toward a knowledge foundation for cognitive neuroscience. Frontiers Neuroinform, 5:17.

[127] Pond, S. L. K., Frost, S. D., and Muse, S. V. (2005). Hyphy: hypothesis testing using phylogenies. Bioinformatics, 21(5):676–679.

[128] Preuschoff, K. and Bossaerts, P. (2007). Adding prediction risk to the theory of reward learning. Annals of the New York Academy of Sciences, 1104(1):135–146.

[129] Quintana, D. S., Rokicki, J., van der Meer, D., Alnæs, D., Kaufmann, T., Córdova-Palomera, A., Dieset, I., Andreassen, O. A., and Westlye, L. T. (2019). Oxytocin pathway gene networks in the human brain. Nature communications, 10(1):668.

[130] Radhakrishnan, R., Matuskey, D., Nabulsi, N., Gaiser, E., Gallezot, J.-D., Henry, S., Planeta, B., Lin, S.-f., Ropchan, J., Huang, Y., et al. (2020). In vivo 5-ht6 and 5-ht2a receptor availability in antipsychotic treated schizophrenia patients vs. unmedicated healthy humans measured with [11c] gsk215083 pet. Psychiatry Research: Neuroimaging, 295:111007.

[131] Radhakrishnan, R., Nabulsi, N., Gaiser, E., Gallezot, J.-D., Henry, S., Planeta, B., Lin, S.-f., Ropchan, J., Williams, W., Morris, E., et al. (2018). Age-related change in 5-ht6 receptor availability in healthy male volunteers measured with 11c-gsk215083 pet. Journal of Nuclear Medicine, 59(9):1445–1450.

[132] Rahim, M., Thirion, B., and Varoquaux, G. (2017). Multi-output predictions from neuroimaging: assessing reduced-rank linear models. In 2017 International Workshop on Pattern Recognition in Neuroimaging (PRNI), pages 1–4. IEEE.

[133] Richiardi, J., Altmann, A., Milazzo, A.-C., Chang, C., Chakravarty, M. M., Banaschewski, T., Barker, G. J., Bokde, A. L., Bromberg, U., Büchel, C., et al. (2015). Correlated gene expression supports synchronous activity in brain networks. Science, 348(6240):1241–1244.

[134] Ripoll-Sánchez, L., Watteyne, J., Sun, H., Fernandez, R., Taylor, S. R., Weinreb, A., Bentley, B. L., Hammarlund, M., Miller, D. M., Hobert, O., et al. (2023). The neuropeptidergic connectome of c. elegans. Neuron, 111(22):3570–3589.

[135] Ritchie, M. E., Phipson, B., Wu, D., Hu, Y., Law, C. W., Shi, W., and Smyth, G. K. (2015). limma powers differential expression analyses for rna-sequencing and microarray studies. Nucleic acids research, 43(7):e47–e47.

[136] Robledo, P., Kaneko, W., and Ehlers, C. (1995). Effects of neurotensin on eeg and event-related potentials in the rat. Psychopharmacology, 118:410–418.

[137] Rokicki, J., Kaufmann, T., De Lange, A.-M. G., van der Meer, D., Bahrami, S., Sartorius, A. M., Haukvik, U. K., Steen, N. E., Schwarz, E., Stein, D. J., et al. (2022). Oxytocin receptor expression patterns in the human brain across development. Neuropsychopharmacology, 47(8):1550–1560.

[138] Romanov, R. A., Tretiakov, E. O., Kastriti, M. E., Zupancic, M., Häring, M., Korchynska, S., Popadin, K., Benevento, M., Rebernik, P., Lallemend, F., et al. (2020). Molecular design of hypothalamus development. Nature, 582(7811):246–252.

[139] Romero-Garcia, R., Whitaker, K. J., Váša, F., Seidlitz, J., Shinn, M., Fonagy, P., Dolan, R. J., Jones, P. B., Goodyer, M., Bullmore, E. T., et al. (2018). Structural covariance networks are coupled to expression of genes enriched in supragranular layers of the human cortex. Neuroimage, 171:256–267.

[140] Rong, J., Haider, A., Jeppesen, T. E., Josephson, L., and Liang, S. H. (2023). Radiochemistry for positron emission tomography. Nature communications, 14(1):3257.

[141] Rozenfeld, E., Tauber, M., Ben-Chaim, Y., and Parnas, M. (2021). Gpcr voltage dependence controls neuronal plasticity and behavior. Nature Communications, 12(1):7252.

[142] Rudy, B., Fishell, G., Lee, S., and Hjerling-Leffler, J. (2011). Three groups of interneurons account for nearly 100% of neocortical gabaergic neurons. Developmental neurobiology, 71(1):45–61.

[143] Russo, S. J. and Nestler, E. J. (2013). The brain reward circuitry in mood disorders. Nature reviews neuroscience, 14(9):609–625.

[144] Sandiego, C. M., Gallezot, J.-D., Lim, K., Ropchan, J., Lin, S.-f., Gao, H., Morris, E. D., and Cosgrove, K. P. (2015). Reference region modeling approaches for amphetamine challenge studies with [11c] flb 457 and pet. Journal of Cerebral Blood Flow & Metabolism, 35(4):623– 629.

[145] Santollo, J. and Eckel, L. A. (2013). Oestradiol decreases melanin-concentrating hormone (mch) and mch receptor expression in the hypothalamus of female rats. Journal of neuroendocrinology, 25(6):570–579.

[146] Sartorius, A. M., Rokicki, J., Birkeland, S., Bettella, F., Barth, C., de Lange, A.-M. G., Haram, M., Shadrin, A., Winterton, A., Steen, N. E., et al. (2024). An evolutionary timeline of the oxytocin signaling pathway. Communications Biology, 7(1):471.

[147] Schaefer, A., Kong, R., Gordon, E. M., Laumann, T. O., Zuo, X.-N., Holmes, A. J., Eickhoff, S. B., and Yeo, B. T. (2018). Local-global parcellation of the human cerebral cortex from intrinsic functional connectivity mri. Cerebral cortex, 28(9):3095–3114.

[148] Scholtens, L. H., Schmidt, R., de Reus, M. A., and van den Heuvel, M. P. (2014). Linking macroscale graph analytical organization to microscale neuroarchitectonics in the macaque connectome. Journal of Neuroscience, 34(36):12192–12205.

[149] Seguin, C., Sporns, O., and Zalesky, A. (2023). Brain network communication: concepts, models and applications. Nature reviews neuroscience, 24(9):557–574.

[150] Shafiei, G., Baillet, S., and Misic, B. (2022). Human electromagnetic and haemodynamic networks systematically converge in unimodal cortex and diverge in transmodal cortex. PLoS biology, 20(8):e3001735.

[151] Shine, J. M., Müller, E. J., Munn, B., Cabral, J., Moran, R. J., and Breakspear, M. (2021). Computational models link cellular mechanisms of neuromodulation to largescale neural dynamics. Nature neuroscience, 24(6):765– 776.

[152] Silva, A. C. and Kim, S.-G. (1999). Pseudo-continuous arterial spin labeling technique for measuring cbf dynamics with high temporal resolution. Magnetic Resonance in Medicine: An Official Journal of the International Society for Magnetic Resonance in Medicine, 42(3):425– 429.

[153] Silva, D. N., Alves, F., Batista, D., and Paulo, O. S. (2018). Trifusion: Streamlining phylogenomic data gathering, processing and visualization. Phylogenomic and population genomic insights on the evolutionary history of Coffee Leaf Rust within the rust fungi, 1001:119.

[154] Singh, K., Cauzzo, S., García-Gomar, M. G., Stauder, M., Vanello, N., Passino, C., and Bianciardi, M. (2022). Functional connectome of arousal and motor brainstem nuclei in living humans by 7 tesla resting-state fmri. NeuroImage, 249:118865.

[155] Sjöstedt, E., Zhong, W., Fagerberg, L., Karlsson, M., Mitsios, N., Adori, C., Oksvold, P., Edfors, F., Limiszewska, A., Hikmet, F., et al. (2020). An atlas of the proteincoding genes in the human, pig, and mouse brain. Science, 367(6482):eaay5947.

[156] Smith, M. D., Wertheim, J. O., Weaver, S., Murrell, B., Scheffler, K., and Kosakovsky Pond, S. L. (2015). Less is more: an adaptive branch-site random effects model for efficient detection of episodic diversifying selection. Molecular biology and evolution, 32(5):1342–1353.

[157] Smith, S. J., Sümbül, U., Graybuck, L. T., Collman, F., Seshamani, S., Gala, R., Gliko, O., Elabbady, L., Miller, J. A., Bakken, T. E., et al. (2019). Single-cell transcriptomic evidence for dense intracortical neuropeptide networks. elife, 8:e47889.

[158] Sperry, R. W. (1963). Chemoaffinity in the orderly growth of nerve fiber patterns and connections. Proceedings of the National Academy of Sciences, 50(4):703–710.

[159] Strand, F. L. (1999). Neuropeptides: regulators of physiological processes. MIT press.

[160] Suárez, L. E., Markello, R. D., Betzel, R. F., and Misic, B. (2020). Linking structure and function in macroscale brain networks. Trends in cognitive sciences, 24(4):302– 315.

[161] Südhof, T. C. (2017). Molecular neuroscience in the 21st century: a personal perspective. Neuron, 96(3):536– 541.

[162] Suyama, M., Torrents, D., and Bork, P. (2006). Pal2nal: robust conversion of protein sequence alignments into the corresponding codon alignments. Nucleic acids research, 34(Suppl_2):W609–W612.

[163] Svensson, E., Apergis-Schoute, J., Burnstock, G., Nusbaum, M. P., Parker, D., and Schiöth, H. B. (2019). General principles of neuronal co-transmission: insights from multiple model systems. Frontiers in neural circuits, 12:117.

[164] Tadel, F., Baillet, S., Mosher, J. C., Pantazis, D., and Leahy, R. M. (2011). Brainstorm: a user-friendly application for meg/eeg analysis. Computational intelligence and neuroscience, 2011.

[165] Talmage, R. V., Grubb, S. A., Norimatsu, H., and VanderWiel, C. J. (1980). Evidence for an important physiological role for calcitonin. Proceedings of the National Academy of Sciences, 77(1):609–613.

[166] Tan, L. L., Oswald, M. J., and Kuner, R. (2021). Neurobiology of brain oscillations in acute and chronic pain. Trends in neurosciences, 44(8):629–642.

[167] Tian, Y., Margulies, D. S., Breakspear, M., and Zalesky, A. (2020). Topographic organization of the human sub-cortex unveiled with functional connectivity gradients. Nature neuroscience, 23(11):1421–1432.

[168] Tóth, A., Záborszky, L., and Détári, L. (2005). Eeg effect of basal forebrain neuropeptide y administration in urethane anaesthetized rats. Brain research bulletin, 66(1):37–42.

[169] Turtonen, O., Saarinen, A., Nummenmaa, L., Tuominen, L., Tikka, M., Armio, R.-L., Hautamäki, A., Laurikainen, H., Raitakari, O., Keltikangas-Järvinen, L., et al. (2021). Adult attachment system links with brain mu opioid receptor availability in vivo. Biological Psychiatry: Cognitive Neuroscience and Neuroimaging, 6(3):360–369.

[170] Uhlén, M., Fagerberg, L., Hallström, B. M., Lindskog, C., Oksvold, P., Mardinoglu, A., Sivertsson, Å., Kampf, C., Sjöstedt, E., Asplund, A., et al. (2015). Tissue-based map of the human proteome. Science, 347(6220):1260419.

[171] Uhlirova, H., Kılıç, K., Tian, P., Thunemann, M., Desjardins, M., Saisan, P. A., Sakadić, S., Ness, T. V., Mateo, C., Cheng, Q., et al. (2016). Cell type specificity of neurovascular coupling in cerebral cortex. elife, 5:e14315.

[172] Uusitalo, M. A. and Ilmoniemi, R. J. (1997). Signal-space projection method for separating meg or eeg into components. Medical and biological engineering and computing, 35:135–140.

[173] van den Heuvel, M. P., Scholtens, L. H., Barrett, L. F., Hilgetag, C. C., and de Reus, M. A. (2015). Bridging cytoarchitectonics and connectomics in human cerebral cortex. Journal of Neuroscience, 35(41):13943–13948.

[174] Van Den Pol, A. N. (2012). Neuropeptide transmission in brain circuits. Neuron, 76(1):98–115.

[175] Van Essen, D. C., Smith, S. M., Barch, D. M., Behrens, T. E., Yacoub, E., Ugurbil, K., Consortium, W.-M. H., et al. (2013). The wu-minn human connectome project: an overview. Neuroimage, 80:62–79.

[176] Van Veen, B. D., Van Drongelen, W., Yuchtman, M., and Suzuki, A. (1997). Localization of brain electrical activity via linearly constrained minimum variance spatial filtering. IEEE Transactions on biomedical engineering, 44(9):867–880.

[177] Vázquez-Rodríguez, B., Suárez, L. E., Markello, R. D., Shafiei, G., Paquola, C., Hagmann, P., Van Den Heuvel, M. P., Bernhardt, B. C., Spreng, R. N., and Misic, B. (2019). Gradients of structure–function tethering across neocortex. Proceedings of the National Academy of Sciences, 116(42):21219–21227.

[178] Vijay, A., Cavallo, D., Goldberg, A., de Laat, B., Nabulsi, N., Huang, Y., Krishnan-Sarin, S., and Morris, E. D. (2018). Pet imaging reveals lower kappa opioid receptor availability in alcoholics but no effect of age. Neuropsychopharmacology, 43(13):2539–2547.

[179] Viñuela, A., Brown, A. A., Buil, A., Tsai, P.-C., Davies, M. N., Bell, J. T., Dermitzakis, E. T., Spector, T. D., and Small, K. S. (2018). Age-dependent changes in mean and variance of gene expression across tissues in a twin cohort. Human molecular genetics, 27(4):732–741.

[180] Wang, H., Qian, T., Zhao, Y., Zhuo, Y., Wu, C., Osakada, T., Chen, P., Chen, Z., Ren, H., Yan, Y., et al. (2023). A tool kit of highly selective and sensitive genetically encoded neuropeptide sensors. Science, 382(6672):eabq8173.

[181] Wei, Y., Scholtens, L. H., Turk, E., and Van Den Heuvel, M. P. (2018). Multiscale examination of cytoarchitectonic similarity and human brain connectivity. Network Neuroscience, 3(1):124–137.

[182] Whitehead, S. D. and Ballard, D. H. (1989). A role for anticipation in reactive systems that learn. In Proceedings of the Sixth International Workshop on Machine Learning, pages 354–357. Elsevier.

[183] Wilkinson, M. and Brown, R. E. (2015). An introduction to neuroendocrinology. Cambridge University Press.

[184] Wise, R. A. and Bozarth, M. A. (1984). Brain reward circuitry: four circuit elements “wired” in apparent series. Brain research bulletin, 12(2):203–208.

[185] Xu, T., Chen, Z., Zhou, X., Wang, L., Zhou, F., Yao, D., Zhou, B., and Becker, B. (2024). The central renin– angiotensin system: A genetic pathway, functional decoding, and selective target engagement characterization in humans. Proceedings of the National Academy of Sciences, 121(8):e2306936121.

[186] Yarkoni, T., Poldrack, R. A., Nichols, T. E., Van Essen, D. C., and Wager, T. D. (2011). Large-scale automated synthesis of human functional neuroimaging data. Nat Meth, 8(8):665.

[187] Zamani Esfahlani, F., Faskowitz, J., Slack, J., Mišić, B., and Betzel, R. F. (2022). Local structure-function relationships in human brain networks across the lifespan. Nature communications, 13(1):2053.

[188] Zambach, S. A., Cai, C., Cederberg Helms, H. C., Hald, B. O., Fordsmann, J. C., Nielsen, R. M., Lønstrup, M., Brodin, B., and Lauritzen, M. J. (2020). Nitric oxide, katp channels and endothelin-1 modulate brain pericyte function, vascular tone and neurovascular coupling. bioRxiv, pages 2020–06.

[189] Zerbino, D. R., Achuthan, P., Akanni, W., Amode, M. R., Barrell, D., Bhai, J., Billis, K., Cummins, C., Gall, A., Girón, C. G., et al. (2018). Ensembl 2018. Nucleic acids research, 46(D1):D754–D761.

[190] Zhang, W., Silva, A. C., Williams, D. S., and Koretsky, A. P. (1995). Nmr measurement of perfusion using arterial spin labeling without saturation of macromolecular spins. Magnetic resonance in medicine, 33(3):370–376.

[191] Zhang, Y., Proenca, R., Maffei, M., Barone, M., Leopold, L., and Friedman, J. M. (1994). Positional cloning of the mouse obese gene and its human homologue. Nature, 372(6505):425–432.

[192] Zhong, W., Barde, S., Mitsios, N., Adori, C., Oksvold, P., Feilitzen, K. v., O’Leary, L., Csiba, L., Hortobágyi, T., Szocsics, P., et al. (2022). The neuropeptide landscape of human prefrontal cortex. Proceedings of the National Academy of Sciences, 119(33):e2123146119.

